# Loss of RNF43/ZNRF3 predisposes to Hepatocellular carcinoma by impairing liver regeneration and altering liver fat metabolism

**DOI:** 10.1101/2020.09.25.313205

**Authors:** Gianmarco Mastrogiovanni, Clare Pacini, Sofia Kakava, Robert Arnes-Benito, Charles R Bradshaw, Susan Davies, Kourosh Saeb-Parsy, Bon-Kyoung Koo, Meritxell Huch

## Abstract

The homologous E3 ubiquitin ligases RNF43/ZNRF3 negatively regulate WNT signalling activation. Recently, both genes have been found mutated in several types of cancers. Specifically, loss-of-function mutations result in adenoma formation in mouse small intestine. However, their role in liver cancer has not been explored yet. Here we describe that hepatocyte-specific deletion of both *Rnf43/Znrf3* results in altered lipid metabolism and a non-alcoholic steatohepatitis (NASH) phenotype in mouse, in the absence of exogenous fat supplementation. The effect is cell-autonomous, as evidenced by the intracellular lipid accumulation detected in mutant liver organoids. Upon chronic liver damage, *Rnf43/Znrf3* deletion results in impaired hepatocyte regeneration, subsequent to an imbalance between hepatocyte differentiation and proliferation, which leads to hepatocellular carcinoma. Remarkably, hepatocellular carcinoma patients with mutations in ZNRF3 also present altered lipid metabolism and poorer survival. Our findings imply that Wnt activation through the RNF43/ZNRF3 module predisposes to liver cancer by altering the liver lipid metabolic ground-state and impairing liver regeneration, which combined, facilitate the progression towards malignancy. Our results highlight the requirement for personalized therapeutic or dietary interventions for those RNF43/ZNRF3 mutated individuals at risk of developing steatosis, NASH and/or liver cancer.

---

The WNT signalling pathway is known for regulating many cellular processes, ranging from cell fate decisions during embryogenesis to maintaining tissue homeostasis in many adult tissues^1^. The key switch in the canonical Wnt pathway is the cytoplasmic protein β-catenin. In the absence of WNT signal, β-catenin is targeted for proteasomal degradation^2^, whereas upon WNT ligand binding to FZDs and LRP5/6, β-catenin accumulates in the cytoplasm and nucleus, where it engages its effectors, the DNA-bound TCF transcription factors^3,4^. In the liver, WNT signalling plays a critical role in tissue regeneration and metabolic zonation as well as in the establishment and progression of liver diseases, including primary liver cancer. Mouse models of liver regeneration, either by tissue resection (partial hepatectomy) or toxic insult (following toxin administration) have demonstrated the importance of proper regulation of the WNT pathway during the regeneration program^5–8^. For instance, ablation of either β-catenin or the LGR4/5-RSPO axis, results in impaired activation and completion of the regenerative process^9–12^. Similarly, the spatial separation of metabolic functions in the liver, known as metabolic zonation, is regulated, in part, by the specific distribution of WNT signals through the perivenous-portal axis. In the central vein (perivenous) region, high levels of WNT signalling emanating from the endothelium^13,14^ regulate lipids and glucose metabolism^15^. In fact, deletion of either β-catenin^11^, TCF4^16^ or LGR4/5^9^ results in reduced expression of zonated pericentral genes (e.g. GS and CYP2E1); conversely, APC liver-specific deletion leads to the expansion of pericentral genes up to the portal area^17^. Also, liver-specific knock-out of TCF4 results in decreased levels of triglycerides, fatty acids and ketone bodies in newborn mice^16^.

The homologous WNT target genes *Rnf43* and *Znrf3*, are two potent negative-feedback regulators of the WNT pathway in Lgr5^+^ intestinal stem cells^18^ and in colon cancer cells carrying activating WNT pathway mutations^19^. Both are E3-ubiquitin ligases that specifically ubiquitinate the cytoplasmic loops of the WNT pathway receptors FZDs^18,19^, which induces their rapid endo-lysosomal degradation. Conversely, R-spondins (RSPO), ligands of LGR4, LGR5 and LGR6 receptors, also interact with RNF43/ZNRF3, which reverses the RNF43/ZNRF3-mediated membrane clearance of FZDs and results in prolonged activation of WNT/FZD/LRP receptor complexes, boosting WNT signal strength and duration^20^.

Inactivating mutations in *Rnf43* and/or *Znrf3* have been observed in a variety of human cancers^73^. We previously reported that intestinal deletion of these two E3 ligases results in enlargement of intestinal crypts and adenoma formation (Koo et al., 2012)^18^, which disappears upon treatment with Porcupine inhibitors (enzyme required for WNT ligand palmitoylation)^21^. Specifically, in primary liver cancer, *Znrf3* is mutated in 3% of hepatocellular carcinomas (HCCs)^22^, while *Rnf43* is mutated in 9.3% of intrahepatic cholangiocarcinoma (iCC)^23^. However, their role in liver cancer, whether drivers or disease risk factors, has not been explored yet. Similarly, ubiquitous (whole-body) deletion of both *Rnf43/Znrf3* results in some liver metabolic changes^9^, however, whether these E3 ligases exert a direct regulation on liver metabolism or whether these result from secondary effects on non-liver tissues has not been elucidated yet.

Here, we aimed at elucidating the specific role of the RNF43/ZNRF3-module in the liver, during homeostasis, hepatocyte regeneration and cancer. We describe that specific deletion of both homologues in adult hepatocytes results in liver degeneration, lipid deposition and altered lipid metabolism under normal diet. The lipid metabolic alteration is cell-autonomous, since *Rnf43*/*Znrf3-null* liver epithelial organoids exhibit significant intracellular lipid accumulation. Furthermore, upon liver damage, hepatocyte specific loss-of-function of both, *Rnf43* and *Znrf3*, results in impaired hepatocyte regeneration, significant liver fibrosis and extensive tissue damage due to impaired hepatocyte differentiation, which progresses to Hepatocellular Carcinoma (HCC). Notably, patients with ZNRF3 mutations present lipid metabolic alterations and poorer prognosis.

Our observations imply that the RNF43/ZNRF3 module does not drive liver cancer directly, instead predisposes to the disease by altering the liver metabolic ground-state which, combined with an impaired ability to regenerate, facilitates the progression towards malignancy.

## Results

### Hepatocyte-specific deletion of *Rnf43/Znrf3* results in liver hyperplasia and foci of lipid deposition

To unveil the role of *Rnf43* and *Znrf3* in the adult liver, we first determined the expression pattern of both genes within the different liver cell populations. We found that both, *Rnf43* and *Znrf3*, were expressed in hepatocytes (**Supplementary Fig. 1a-b**), in agreement with a previous report^9^. In addition, we also detected expression for both genes in EpCAM^+^ ductal cells, with *Rnf43* being predominantly expressed in hepatocytes and *Znrf3* in ductal cells. We did not detect expression of neither of the two genes in CD31^+^ endothelial cells; whereas *Znrf3* was slightly expressed in macrophages (CD11b^+^) (**Supplementary Fig. 1a-b**).

Hence, to study the hepatocyte-specific role of *Rnf43/Znrf3*, we next generated a hepatocyte-specific compound mutant mouse, by taking advantage of our previously reported conditional alleles for *Rnf43/Znrf3*^*flox*18^, which allow specific deletion of the two genes simultaneously (**Supplementary Fig. 1c**). We generated hepatocyte-specific *Rnf43/Znrf3* mutant mice by either crossing *Rnf43/Znrf3*^*flox*^ mutant mice with a tamoxifen-inducible *AlbCre-ERT2* reporter^24^, to generate the compound mice *AlbCre-ERT2/Rnf43-Znrf3*^*flox*^, or by injecting *Rnf43/Znrf3*^*flox*^ mice with an hepatotropic AAV8-TGB-Cre virus (see methods) (**Fig. 1a**), hence generating *Rnf43/Znrf3* hepatocyte-specific mutant mice (*Rnf43/Znrf3*^*del*^ mice, from hereon). Using either recombination approach, we observed hepatocyte-specific deletion of the two genes at all time points analysed (from 1 week up to 7 months post induction) (**Supplementary Fig. 1d**). No recombination was detected in non-tamoxifen treated mice (**Supplementary Fig. 1e**).

**Figure 1.**
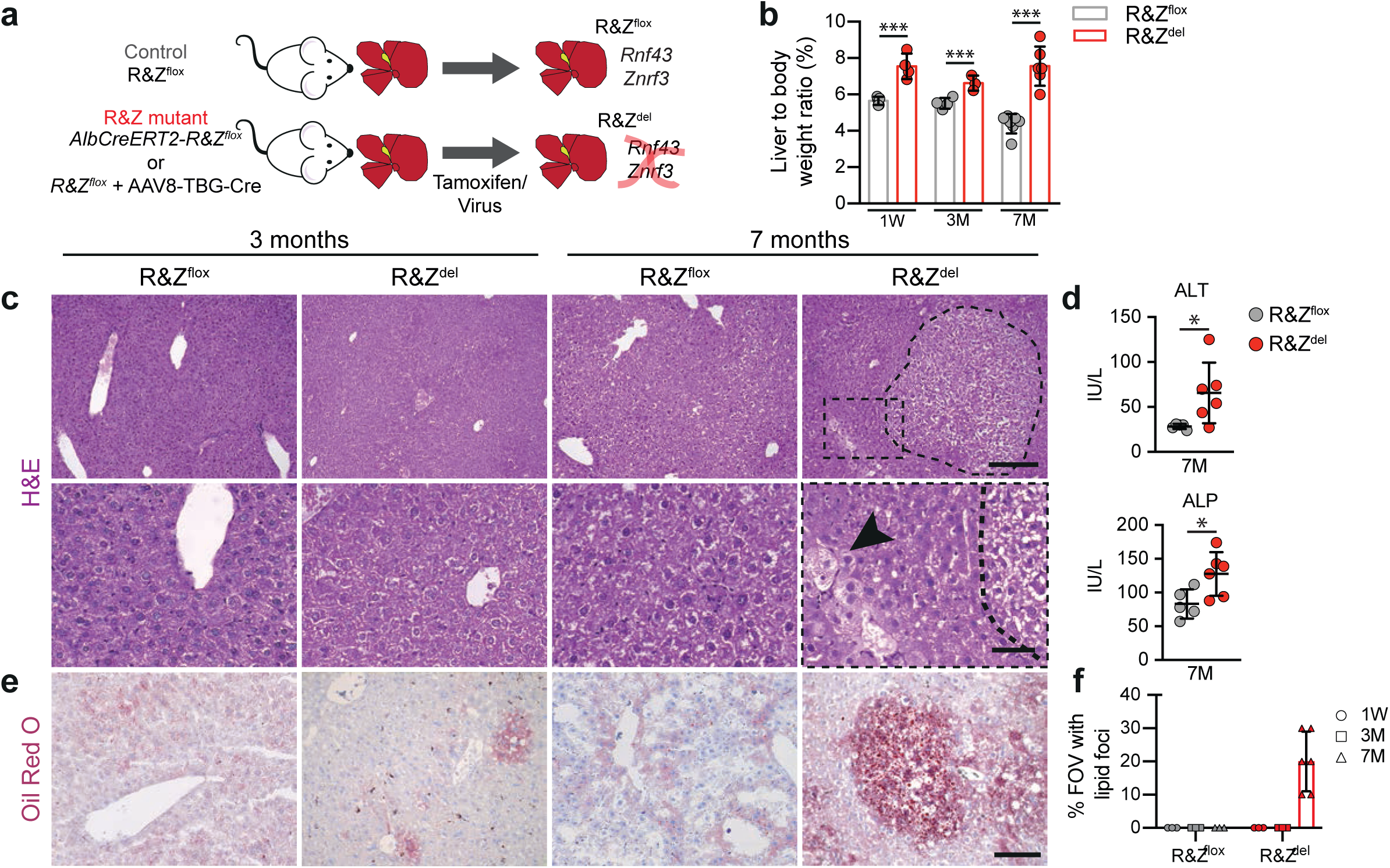
Liver-specific *Rnf43 & Znrf3* (R&Z) deletion induces hepatomegaly, tissue degeneration and lipid foci. Liver-specific *Rnf43 & Znrf3* (R&Z) mutant mice were obtained by using R&Z floxed mice (R&Z^flox^) and then either cross them with AlbCreERT2 mice and injected with tamoxifen, or inject them with AAV8-TBG-Cre. Livers were collected 1 week and 3 and 7 months later and processed for histopathological analysis. R&Z^flox^ mice lacking the AlbCreERT2 allele were also injected with tamoxifen or AAV8-Null virus and used as controls. **a)** Experimental design. **b)** R&Z liver specific deletion results in hepatomegaly. Graph represents the % of liver-to-body weight ratio. Results are presented as mean +/- SD from R&Z^flox^-1W, n=5 mice; R&Z^del^-1W and R&Z^flox^-3M, n=4 mice; R&Z^del^-3M, n=3 mice; R&Z^del^-7M, n=6 mice; R&Z^flox^-7M, n= 7 mice. Two-tail t-test was used; ***P<0.001. **c)** Histopathological analysis revealed the presence of nodules with white changes and cellular degeneration, as evidenced by the presence of ballooning hepatocytes (bottom left panel) in the livers of *Rnf43* & *Znrf3* (R&Z) mutant mice compared to littermate controls. Representative pictures are shown. Scale bar is 50μm for top panel and 100μm for bottom panel. Arrowhead, ballooning degeneration. **d)** Graphs show serum levels for the serum markers ALT and ALP. Analysis was performed 7 months after deletion. Data represent mean +/- SD of n=5 R&Z^flox^ and n=6 R&Z^del^ samples. Two-tail t-test was used; *P<0.05. **e-f)** Oil red O staining revealed the presence of liver fat accumulation as lipid foci. Representative pictures are shown **(e)**. Scale bar, 100μm. Graph represents % of FOV with or without lipid foci **(f)**.

To assess the effect of hepatocyte-specific loss-of-function of *Rnf43/Znrf3*, we collected *Rnf43/Znrf3*^*del*^ mutant livers at 1 week, 3 months and 7 months after deletion (**Supplementary Fig. 2a**). *Rnf43/Znrf3*^*del*^ mutant mice exhibited significant hyperproliferative hepatomegaly already 7 days after the deletion (**Fig. 1b** and **Supplementary Fig. 2h**, left panel), resembling the effects of hepatocyte-specific *Apc* mutant mice^17^. Notably, the hyperplasia and hepatomegaly were maintained throughout the course of the study, with a pronounced increase in the number of proliferating hepatocytes (Ki67^+^) over time, all over the liver parenchyma (**Supplementary Fig. 2d-e,h)**. *Rnf43/Znrf3*^*del*^ mutant livers also presented a marked increase over time on the Wnt/β-catenin liver zonated gene GS, compared to control littermates, which invaded up to 50% of the tissue (**Supplementary Fig. 2f-h**). Notably, at 7 months, *Rnf43/Znrf3*^*del*^ mice exhibited a profound cellular degeneration, with clear cell nodules formed by hepatocytes with nuclear alterations (crenation and karyorrhexis) and ballooned and degenerative hepatocytes present throughout the parenchyma (**Fig. 1c**, right panel and **Supplementary Fig. 2b**, left panel). These results were in agreement with the biochemistry analysis of the mice blood, which revealed a significant elevation of the liver parameters AST and ALP (**Fig. 1d**). Tissue degeneration was specifically observed following long-term loss-of-function, while at 1 week and 3 month, *Rnf43/Znrf3*^*del*^ mice did not present any apparent histo-morphological changes when compared to littermate controls (**Fig. 1c**, left panel and **Supplementary Fig. 2h**, R&Z^flox^ panels). Also, individual homozygous mutants did not yield any discernible phenotype (**Supplementary Fig. 2c,h**).

Tissue degeneration and ballooned hepatocytes are observed in a variety of acute and chronic liver diseases^25,26^, but specifically, hepatocyte ballooning degeneration is considered a hallmark of chronic lipotoxic damage and steatohepatitis/NASH (inflamed fatty liver)^25,27,28^. Notably, histopathological analysis of our *Rnf43/Znrf3*^*del*^ mutant livers indicated that at 7 months after deletion, all mutant livers exhibited features of steatohepatitis, which was confirmed by the presence of mild inflammation (**Supplementary Fig. 2b**, right panel) and foci of fat deposition, as determined by Oil Red-O staining, which allows the visualization of neutral triglycerides and lipids (**Fig. 1e-f**). We did not observe any foci of fat accumulation in any of the WT littermates analysed.

Collectively, these data suggested that *Rnf43/Znrf3*^*del*^ might play a role in maintaining adult liver tissue homeostasis and liver lipid metabolic rheostat.

### *Rnf43/Znrf3* regulate liver fat metabolism and lipid accumulation cell-autonomously

To gain a detailed resolution of the role of *Rnf43*/*Znrf3* in hepatocyte maintenance and metabolism, we next performed global gene expression analysis (RNAseq) of *Rnf43/Znrf3*^*del*^ and corresponding littermate control livers at the time point where fat deposition foci were detected (7 months after gene deletion). PCA analysis revealed that *Rnf43/Znrf3*^*del*^ mutants clustered together and separated from control littermates (**Supplementary Fig. 3a**, squares). We found 402 genes differentially expressed (FDR<10% and FC>1) in *Rnf43/Znrf3*^*del*^ livers compared to controls (**Supplementary Fig. 3b**, first row). Notably, gene ontology (GO) analysis revealed that a significant proportion of the upregulated genes were involved in lipid/phospholipid metabolism, being “lipid metabolic process” the top most significantly enriched category (**Fig. 2a** and **Supplementary Data Set 1**). Indeed, when examining the differentially expressed genes we found that 18.4% of them (74 out of 402) were lipid metabolic genes (**Fig. 2b**). Within these, we found genes encoding for enzymes involved in the biosynthesis of triglycerides, fatty acids and phospholipids, such as *Fads2* (fatty acids), *Cyp7a1* (cholesterol) and *Gpat4* (phospholipids), as well as enzymes involved in diacylglycerol (precursors of triacylglycerol and phospholipids) biosynthesis such as *Mogat2, Mogat1, Lpin2 and Lpin1*, among others (**Fig. 2c,d** and **Supplementary Fig. 3c**, right panel). In addition, genes involved in lipid uptake (*Cd36*), adipogenesis (*Cidea, Pparg*) and lipid transfer (*Pltp*) were also significantly upregulated (**Fig. 2c,d** and **Supplementary Fig. 3c**, right panel). Conversely, genes involved in phospholipid oxidation (*Pla2g7*) and cholesterol hydroxylation (*Hsd11B1*) were within the most downregulated genes (**Fig. 2d**). These molecular changes in metabolic genes were also observed at 3 months after deletion (**Supplementary Fig. 3c**, left panel), albeit at this earlier time point we had not detected a significant increase in fat accumulation at the histological level (**Fig. 1e-f**).

**Figure 2.**
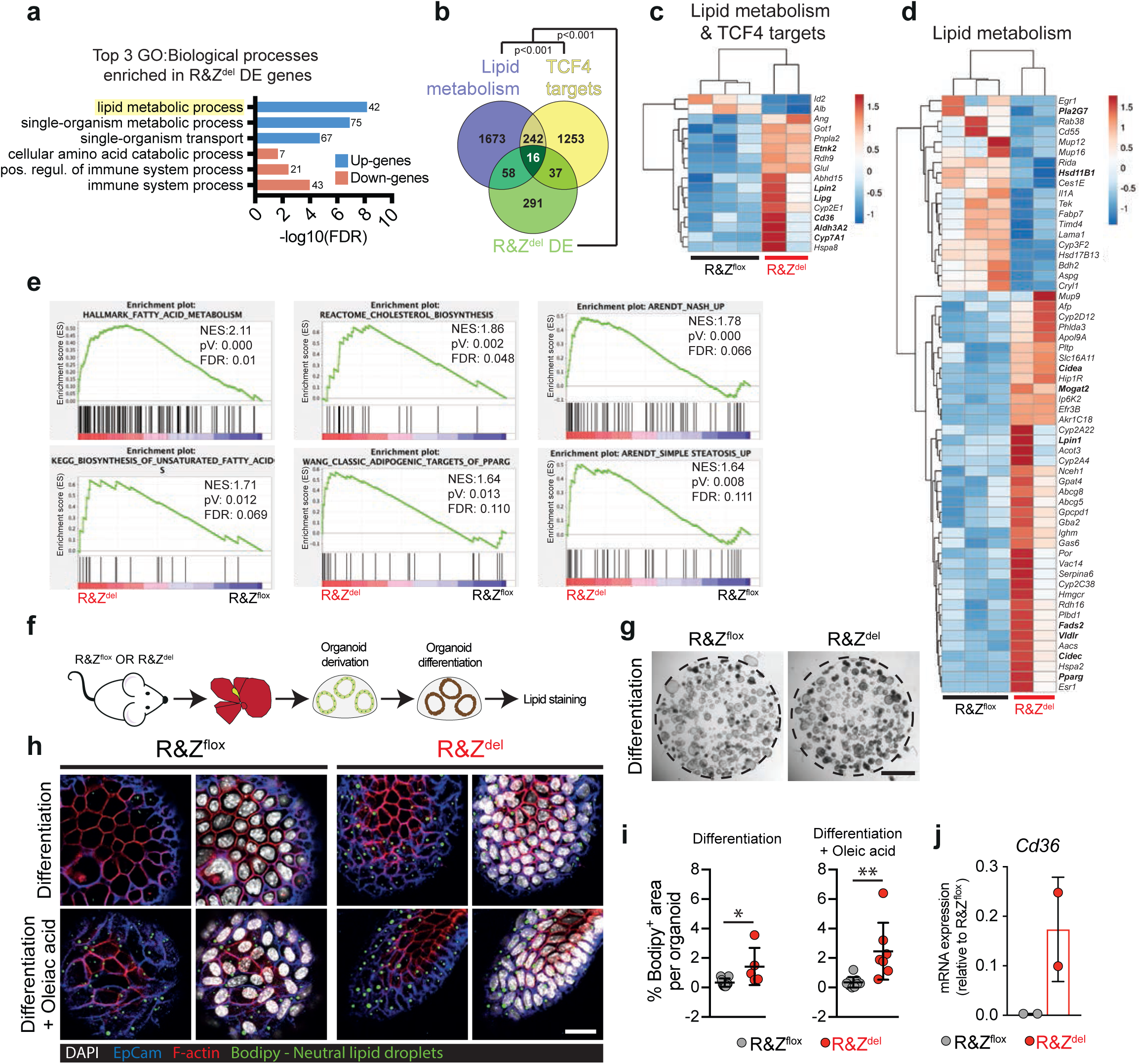
Constitutive activation of the WNT canonical pathway in *Rnf43 & Znrf3* (R&Z) null hepatocytes results in lipid metabolic changes and a steatohepatitis (NASH) phenotype due to a cell-autonomous accumulation of lipid droplets in hepatic cells. **a-e)** Liver tissues from 7 months old R&Z^del^ mice (n=2) and R&Z^flox^ littermates (n=3) were collected and processed for RNAseq analysis. Differential Expressed (DE) gene profiles were generated between the 2 groups. **a)** Graph showing top 3 gene ontology (GO) terms significantly enriched for genes up- (blue) and down- (pink) regulated in R&Z^del^ compared to R&Z^flox^. The numbers denote the number of genes associated to each term. **b)** Venn diagram showing correlation between genes involved in lipid metabolism (purple), TCF4 target genes (yellow) and genes differentially expressed in R&Z^del^ livers (green). Lipid metabolism genes were selected from the following genesets from mSigDB: Hallmark fatty acid metabolism; Reactome cholesterol biosynthesis; Wang adipogenic targets of PPARG; and the GO term GO:0006629 lipid metabolic process (including children terms).TCF4 targets were obtained from Boj et al. Cell. 2012 ChIP-seq data by selecting genes with a TCF4 binding site +/- 5kb from TSS. The numbers denote the number of overlapping genes between DE genes (R&Z^del^ vs R&Z^flox^) and lipid metabolism genes and TCF4 targets. Statistically significant overlap was observed when comparing the overlap to random ChIP profiles. Details are given in **Supplementary Data Sets 2. c-d)** Heat-map analysis of the RPKM values (raw z-scored) of the genes overlapped with lipid metabolism and TCF4 targets **(c)** or lipid genes only **(d). e)** GSEA revealed that 31 out of 211 gene sets involved in lipid metabolism were significantly enriched in our DE gene signatures (FDR<25%). Representative plots are shown. Details of all the positively and negatively correlated signatures are given in **Supplementary Data Sets 1. f-j)** *Rnf43* & *Znrf3* (R&Z) mutant organoids revealed that the lipid metabolic changes observed at the transcriptomic level are caused, in part, by a cell-autonomous accumulation of lipid droplets in hepatic cells. **f)** Schematic representation of the experiment. **g)** Brightfield pictures of liver organoids after culturing in differentiation medium for 10 days. Scale bar, 2mm. **h)** Mouse liver organoids derived from either R&Z^flox^ or R&Z^del^ livers were stained with Bodipy during expansion or after differentiation into hepatocytes. Scale bar is 30μm. **i)** Graphs showing bodipy staining quantification. Two-tail t-test was used; *p<0.05; **p<0.01. Data represents mean +/- SD of a minimum of 5 organoids per condition. **j)** RT-qPCR expression analysis of the lipid transporter Cd36. Data represents mean +/- SD of 2 biological replicates.

Subsequent Gene Set Enrichment Analysis (GSEA) against a list of all curated metabolic dataset confirmed a positive enrichment for metabolic datasets (31 out of 448 datasets) including fatty acid metabolism (*Cidea, Glul, Cd36, Cpt1a, Ppara, Fasn*), cholesterol biosynthesis (*Mvd, Hmgcr*) and PPARG targets (*Cidec, Glul, Cd36, Pnpla2*), among others (**Fig. 2e, Supplementary Data Set 1** and **Supplementary Fig. 3d**). Notably, we also detected positive enrichment against a dataset containing genes upregulated in steatotic and NASH patients^29^, in agreement with our histological findings and confirming, at the molecular level, the steatotic/NASH profile of *Rnf43/Znrf3*^*del*^ mice.

To ascertain whether the changes observed were due to the hyper-activation of canonical WNT pathway we compared the genes differentially expressed to a publish list of liver TCF4 targets obtained by ChIP-seq^16^. As expected from negative regulators of the pathway, we found that 13.2% of our DE genes (53 of 402 genes, p<0.001) were canonical liver WNT/TCF4 targets, such as *Axin2, Sp5, Cyp2e1, Tnfrsf19* and *Rnase4* (**Fig. 2b** and **Supplementary Fig. 3e**), while pericentral and periportal genes were inverted compared to wild-type mice (**Supplementary Fig. 3f**), confirming the reported role of canonical Wnt signalling in the maintenance of liver zonation^9,16^. Interestingly, when overlapping the list of all our DE genes that were lipid metabolic genes with the TCF4 targets we found that 21.6% of these were also *bona-fide* canonical Wnt targets (16 out of 74) (p<0.001) (**Fig. 2b-c**). These included enzymes involved in lipid, cholesterol, triglyceride and phospholipid biosynthesis such as *Aldh3a2, Cyp7a1, Lpin2* and *Etnk2* respectively, as well as lipid and lipid-precursor import genes like *Cd36*^30^ and *Lipg*^31^, respectively (**Fig. 2c**).

To delineate whether the lipid alterations observed were due to a cell-autonomous effect of *Rnf43/Znrf3* in hepatocyte metabolism, or instead secondary to the overall changes that occurred to the tissue overtime, we studied the impact of deleting these two E3 ligases in hepatic cells *ex-vivo*. For that, we took advantage of our liver organoid culture system, that enables the expansion of mutant epithelial cells derived from mouse tissue and the subsequent generation of hepatocyte-like cells following differentiation, in the absence of a mesenchymal/endothelial niche^32^. Hence, we generated *Rnf43/Znrf3*^*del*^ liver organoids (see methods) and, upon differentiation, we analysed their lipid content by staining for neutral lipids (**Fig. 2f-j**). We observed a significant increase in the number of lipid droplets in *Rnf43/Znrf3* mutant organoids, effect that was even more profound when we supplemented the cultures with free-fatty acids (oleic acid) (**Fig. 2h-i**). This increase correlated with the upregulation of the free-fatty acid transporter *Cd36* (**Fig. 2j**). These results suggested that *Rnf43/Znrf3*-null hepatocytes might overproduce, as well as over uptake, and, consequently accumulate, fatty acids in a cell-autonomous manner (**Fig. 2h-j**).

Taken together, we identified a cell-autonomous role for RNF43/ZNRF3 as potential gatekeepers of the lipid metabolic ground-state in hepatocytes, in part through the regulation of canonical WNT signalling.

### *Rnf43/Znrf3* mutant mice exhibit extensive fibrosis and impaired regeneration upon damage due to incomplete hepatocyte maturation

Homeostatic liver cellular turnover is slow, yet the liver displays a great regenerative capacity even upon significant hepatocyte damage. This regenerative ability, though, is not unlimited and chronic liver diseases, including liver cancer, are often preceded by damage to the parenchyma^26^. Our observations that *Rnf43/Znrf3* livers exhibit significant degree of parenchymal degeneration (**Fig. 1**) and a steatohepatitis phenotype that correlates with a NASH/cancer signature (**Fig. 1-2**), prompted us to hypothesise that the homeostatic turnover of the hepatic parenchyma might be impaired in *Rnf43/Znrf3*^*del*^ mice. Hence, we decided to investigate the role of RNF43/ZNRF3 during damage-regenerative response. For that, we challenged *Rnf43/Znrf3*^*del*^ mice with repeated doses of carbon tetrachloride (CCl4) (twice a week for a total of 6 weeks)^33^ in order to induce chronic liver damage (**Fig. 3a**). To prevent regeneration from un-recombined hepatocytes, tamoxifen was also re-injected (11, 30 and 35 days before the end of the chronic damage protocol), which ensured a >80% recombination of both alleles in all mice, at all time points analysed (**Fig. 3a** and **Supplementary Fig. 4a**).

**Figure 3.**
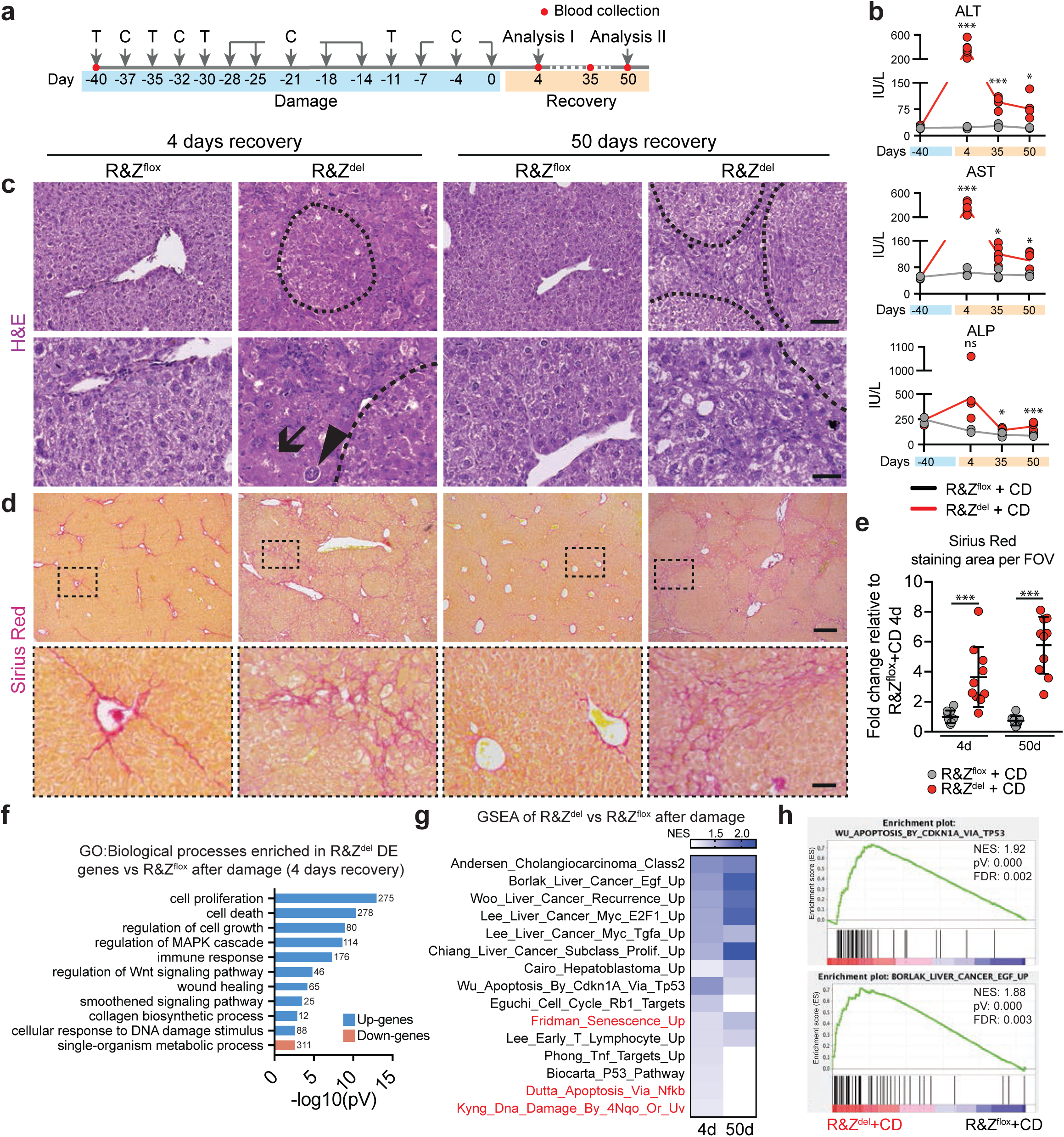
Liver-specific *Rnf43 and Znrf3* (R&Z) deletion leads to improper tissue regeneration after chronic CCl4 treatment due to increased damage. Liver-specific R&Z mutant mice were injected with CCl4 for 6 weeks to induce chronic damage to the liver and tissues were collected 4 and 50 days later and processed for histopathological analysis. Blood was also collected at different time points to assess liver damage (red dots). R&Z^flox^ mice were also injected with tamoxifen and CCl4 and used as controls. **a)** Timeline of the experiment. T, tamoxifen; C, CCl4. **b)** Graphs showing levels of the serum markers ALT, AST and ALP. Analysis was performed at time −40 days (before damage) and at 4, 34 and 50 days of recovery. R&Z^flox^, n=3; R&Z^del^, n=3 at time 0 and n=5 at 4, 34 and 50 days recovery. Lines connect mean between time points. Two-tail t-test was used; *P<0.05; **P<0.01; ***P<0.001; ns, not significant. **c)** Representative pictures of H&E staining. Histopathological analysis revealed increased tissue damage as evidenced by the presence of giant hepatocytes, de-cellularised areas and regenerative nodules in the livers of Rnf43 & Znrf3 mutant mice compared to littermate controls. Scale bar, 100μm for top panel and 50μm for bottom panel. Arrow, de-cellularised area; arrowheads, giant hepatocytes; dashed line, regenerative nodules. **d-e)** Mutant livers after chronic damage present mild fibrosis as shown by increased Pico-Sirius red staining. Representative pictures are shown **(d)**. Scale bar is 500μm. Graphs represent fold change increase of Pico-Sirius red staining area **(e)**. Data represent mean +/- SD of n=2 mice per group, 5 fields per mouse. Data represent mean ± SD; ***P<0.001; two-tail t-test was used. **f)** Graph showing selected gene ontology (GO) terms significantly enriched for genes up-(blue) and down-(pink) regulated in R&Z^del^ compared to R&Z^flox^ after damage. Numbers on the bars denote number of genes associated to each term. Full list can be found in **Supplementary Data Sets 1. g)** Heat map of the normalised enrichment score (NES) of selected GSEA datasets. **h)** Representative GSEA plots for R&Z^del^ livers 4 days after recovery.

As a proxy for liver damage, we first quantified the levels of transaminases in serum before the first dose of CCl4 (day -40) and at several time points after the last dose (day 4, 35 and 50 of recovery). We observed significantly higher levels of ALT, AST and ALP in our mutant mice compared to WT littermates undergoing the same damage regimen (**Fig. 3b**). This increase on the transaminases levels was persistent through all the time points analysed, even after > 2 months of recovery, suggesting that the *Rnf43/Znrf3*^*del*^ livers had an impaired ability to regenerate after a chronic insult (**Fig. 3b**). In line with these results, histological analysis revealed that *Rnf43/Znrf3*^*del*^ livers exhibited extensive tissue damage, with decellularized/necrotic regions, presence of giant hepatocytes and marked immunological infiltrate, all histological hallmarks of prominent tissue damage (**Fig. 3c**). In addition, mutant livers were significantly bigger (**Supplementary Fig. 4b**) and presented multiple regenerative and pre-neoplastic nodules visible at the macroscopic examination (**Supplementary Fig. 4c**), which were composed of hepatocytes with decreased nuclear-to-cytoplasmic ratio. In addition, the tissue presented significant fibrosis and extensive ductular reaction, as evidenced by a significant increase in collagen deposition (Sirus Red area; **Fig. 3d-e**) and the number of pan-cytokeratin (PCK) positive cells (**Supplementary Fig. 4d-e**), respectively. Similarly, HMGB1 staining (marker of cellular injury when not nuclear) confirmed that *Rnf43/Znrf3*^*del*^ livers were extensively damaged (**Supplementary Fig. 4f-g**). Notably, these lesions persisted up to the last time point analysed (50 days recovery, i.e., 50 days after the last CCl4 injection). Expectedly, we also found a significant increase in the GS staining area after 50 days of recovery, but not after 4 days, when the cells usually expressing this marker are highly damaged by the CCl4 (**Supplementary Fig. 4h-i**). None of these serum and histological findings were detected in control littermates, which had mostly recovered already at day 4 after the last CCl4 injection (**Fig. 3b-d** and **Supplementary Fig. 4c-g**), neither in mutant livers not exposed to damage (see **Fig. 1** and data not shown). Importantly, the combination of these histological features (fibrosis, tissue damage, regenerative/pre-neoplastic nodules and ductular reaction) defines human liver cirrhosis^34^ and indicated that the *Rnf43/Znrf3*^*del*^ livers were still undergoing a damage-regenerative response at day 50 of recovery, when the WT tissue had fully regenerated.

Wnt ligands produced by macrophages^35^, endothelial cells^13,14^ and hepatocytes^74^ are essential to activate the regenerative response of hepatocytes to injury. Our results, though, seemed in an apparent discrepancy with the well-documented role of active Wnt signalling in regeneration^6^. However, detailed analysis indicated that the tissue did not fail in activating the regeneration response, as we found increased number of proliferative (Ki67+) cells and significant hepatomegaly upon damage (**Supplementary Fig. 4b, j-k**), in agreement with the reports of active β-catenin^36^ and liver-specific APC mutant mice^37^. Hence, to gain further insights in the response to damage, we performed RNAseq analysis of the CCl4-damaged *Rnf43/Znrf3*^*del*^ livers at days 4 and 50 of recovery and compared the expression patterns to WT control littermates subjected to the same CCl4 regimen. PCA analysis revealed that PC1 component separated damage *vs* undamaged, while the PC3 separated both genotypes, the latter in agreement with the *Rnf43/Znrf3*^*del*^ livers under homeostasis (**Supplementary Fig. 3a**). We found a significant number of genes differentially expressed (2989 at day 4 and 1438 at day 50) between mutant and WT livers indicating a significantly different response to damage between both genotypes (**Supplementary Fig. 3b** and **Supplementary Data Set 1**). Interestingly, we found several metalloproteases involved in tissue remodelling (*Mmp7, Mmp11, Mmp2, Mmp23*) and components of the caspase-mediated apoptosis signalling pathway (*Casp12, Pycard*) amongst the top upregulated genes. GO term analysis of the significantly upregulated genes indicated that many biological processes involved in regeneration and proliferation were activated, with “regulation of cell growth”, “proliferation” and “wound healing” and the signalling pathways Wnt, Shh and MAPK amongst the top significantly enriched GO terms (**Fig. 3f**). Interestingly, GSEA analysis revealed a significant enrichment for liver cancer datasets, including cholangiocarcinoma and hepatoblastoma, among others, as well as datasets associated with tissue damage, including apoptosis, senescence, DNA damage and inflammation (**Fig. 3g-h**). These molecular signature changes were detected not only after 4 days of recovery but also at the 50 days recovery time point. These results confirming that, at the molecular level, *Rnf43/Znrf3*^*del*^ livers have not completed the regenerative response, even 2 months after the last damage dose, in line with our histological findings.

Next, we asked whether the impaired regeneration could be explained by the significant activation of the Wnt pathway. We found that canonical Wnt signalling is dynamically regulated in WT mice during the damage response upon CCl4 damage, with canonical Wnt becoming upregulated as early as day 3 after CCl4 [in agreement with our previous report^32^] and its levels quickly returning to homeostasis once the tissue completes the proliferative phase and starts differentiating, between day 13 and 42 after damage (**Supplementary Fig. 5a**). Hence, we hypothesized that the persistent hyperactivation of the pathway might maintain the hepatocytes in a constant proliferative state that would hinder their complete maturation and, consequently, impair the tissue to finalize the regenerative process. To investigate that, we performed GSEA between the damaged *Rnf43/Znrf3*^*del*^ mice and signatures of both mature hepatocytes and embryonic hepatoblasts (embryonic liver progenitors)^38,39^ (**Fig. 4a**). Interestingly, we observed a significant negative enrichment against hepatocyte signatures. Several mature hepatocyte markers including genes involved in the complement complex (*C9, C8a, C8b, CP*), basic hepatocyte functions (*AR, Baat, Tat*) as well as well-known transcription factors important for hepatocyte-fate specification during embryonic liver development (*Prox1, Hnf4a*) were significantly downregulated compared to wild-type littermates under the same damage regimen (**Fig. 4b** and **Supplementary Fig. 5b**). Conversely, we also observed a positive enrichment for hepatoblast signatures, with bonafide hepatoblast markers such as *Afp* and *Krt8* being significantly upregulated (**Fig. 4a-b**). Similar trend was observed in the mutant compared to WT mice in homeostasis (**Fig. 4a**, top panel and **Supplementary Fig. 5c**).

**Figure 4.**
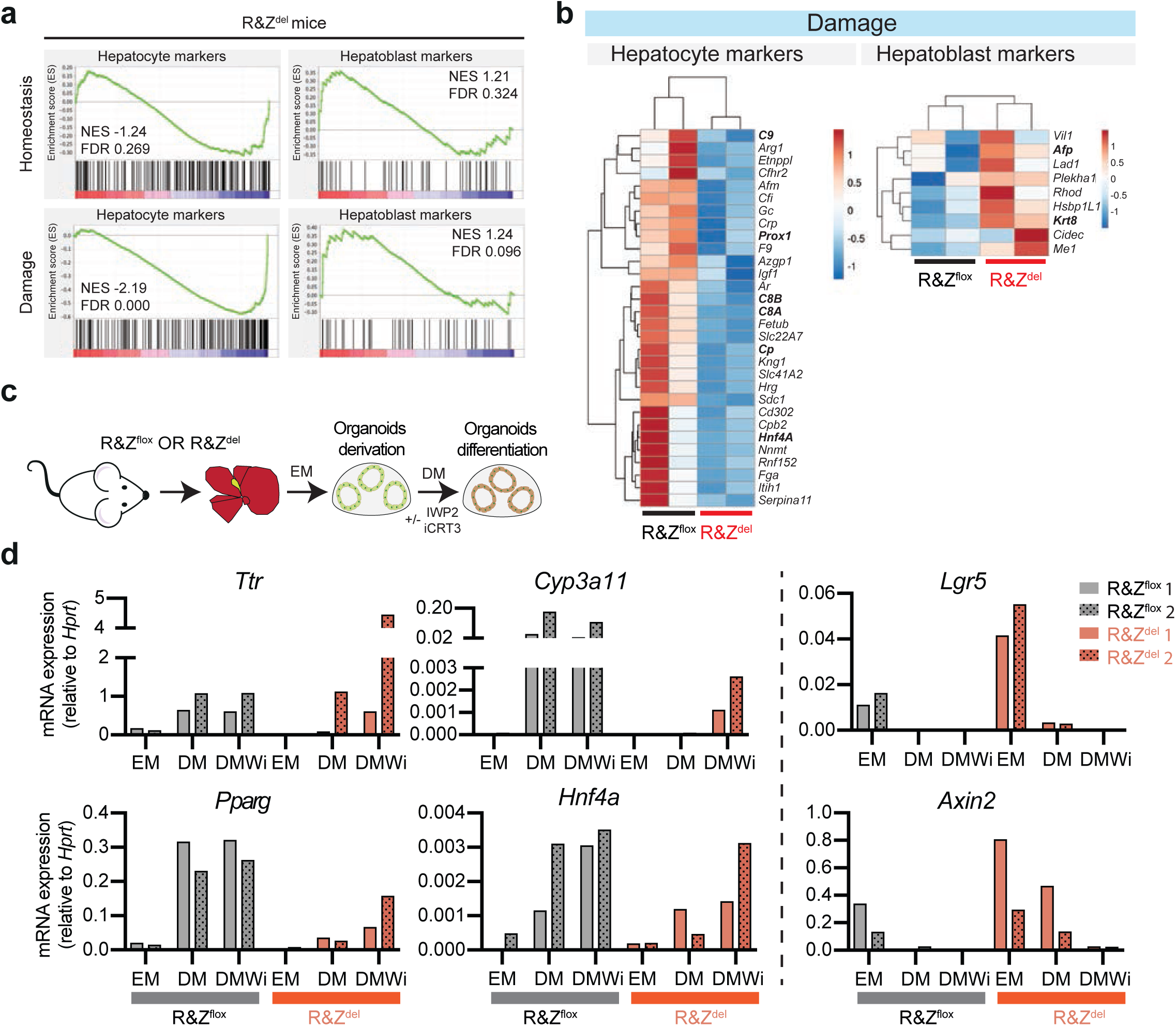
*Rnf43* and *Znrf3* (R&Z) deletion results in impaired maturation of hepatocytes. **a)** GSEA analysis revealed significant negative correlation between hepatocytes signature and damaged R&Z^del^ livers. Plots for hepatocytes and hepatoblats signatures are shown, for both damaged and undamaged livers. **b)** Heat-map analysis of the RPKM values (raw z-scored) showing hepatocyte and hepatoblast genes enriched in the GSEA signature. c-e) Rnf43 & Znrf3 mutant organoids fail to differentiate into hepatocytes due to the activation of the WNT pathway. **c-e)** Organoids were isolated from either R&Z^flox^ or R&Z^del^ livers, and subsequently cultured in expansion medium (EM) or differentiated into hepatocytes using our standard differentiation medium (DM) or this differentiation medium supplemented with small molecule inhibitors of the WNT pathway (DMWi) (IWP2 and iCRT3) and analyzed for the expression of mature/differentiation markers. **c)** Schematic of the experiment. **d)** qPCR expression analysis of organoid from R&Z^flox^ (n=2) and R&Z^del^ (n=2) livers for the hepatocyte differentiation genes *Ttr, Cyp3a11, Pparg* and *Hnf4a* and of the WNT target genes *Lgr5* and *Axin2*. Graphs present the values of each biological replicate normalized by the value of the house-keeping gene *Hprt*.

To determine whether the reduced regeneration and differentiation capacity of the *Rnf43/Znrf3*^*del*^ mice was due to a cell-autonomous effect of the WNT hyperactivation in hepatocytes, we took advantage of our organoid culture system and differentiated *Rnf43/Znrf3*^*del*^ mouse liver organoids into hepatocyte-like cells using our published differentiation protocol (DM)^32^ or upon supplementing this with IWP2^40^ and iCRT3^41^ (DMWi), inhibitors of WNTs secretion and of β-catenin/TCF4 interaction, respectively, to completely inactivate the WNT pathway (**Fig. 4c** and **Supplementary Fig. 5d**). In agreement with our mouse data, *Rnf43/Znrf3*^*del*^ organoids exhibited less differentiation capacity than control WT organoids, as assessed by qPCR analysis of common hepatocyte markers such as *Ttr, Cyp3a11, Pparg* and *Hnf4a*. Conversely, stem cell and WNT target genes like *Lgr5* and *Axin2* were significantly upregulated (**Fig. 4d**). Interestingly, blockade of WNT (produced by the organoid epithelial cells) by IWP2 combined with inhibition of the pathway downstream of the RNF43/ZNRF3 receptors by iCRT3 (DMWi medium) resulted in partial rescue of the differentiation ability of the cells, with significantly higher levels of *Ttr, Cyp3a11, Pparg* and *Hnf4a* when compared to DM only. DMWi treatment abolished the expression of the Wnt targets *Lgr5* and *Axin2*, as expected (**Fig. 4d**).

Collectively, these results imply that the hyperactivation of the WNT pathway impairs the ability of *Rnf43/Znrf3*^*del*^ hepatocytes to terminate the differentiation program, which results in a reduced regenerative capacity and a persistent tissue damage that leads to liver cirrhosis upon chronic injury.

### RNF43/ZNRF3 deletion predisposes to hepatocellular carcinoma

Fatty liver (steatosis) and non-alcoholic steatohepatitis (NASH) are predisposing risk factors for liver cirrhosis and subsequent development of hepatocellular carcinoma (HCC)^26,27^. To assess whether the NASH and cirrhotic phenotypes observed in *Rnf43/Znrf3*^*del*^ mice would progress into a malignant hepatocellular carcinoma stage we subjected the mice to chronic injury and collected the livers ∼5 months later (170 days after the last dose of CCl4) (**Fig. 5a**). We noted the appearance of tumoral nodules of different sizes already at the macroscopic examination of the tissue (**Supplementary Fig. 6a-b**). Histopathological analysis confirmed that these were HCC (HCC) or HCC with ductal features (CHC, mixed subtype), with a penetrance of 66.7% (**Fig. 5b-c** and **Supplementary Fig. 6b-c**). Both types of lesions (HCC and CHC) were characterised by small cells that presented a compact pattern without trabecular formation and 2-3 cell thick plates, fat accumulation and a significant number of proliferative (Ki67^+^) cells and neo-vascularization, as evidenced by CD34 staining (**Fig. 5c** and **Supplementary Fig. 6b**). The mixed HCC (CHC) lesions in addition presented glandular structures formed by PCK^+^ ductal cells (**Supplementary Fig. 6b**). None of these malignant features were observed in any of the damaged-WT (**Fig. 5b** and **Supplementary Fig. 6c**) or undamaged mutant (**Fig. 1**) mice. As expected, the non-tumorous tissue also presented foci of fat accumulation and increased fibrosis, proliferation and ductular reaction as assessed by Oil red-O, Sirius red, Ki67 and PCK staining, respectively (**Supplementary Fig. 7a,c-d**).

**Figure 5.**
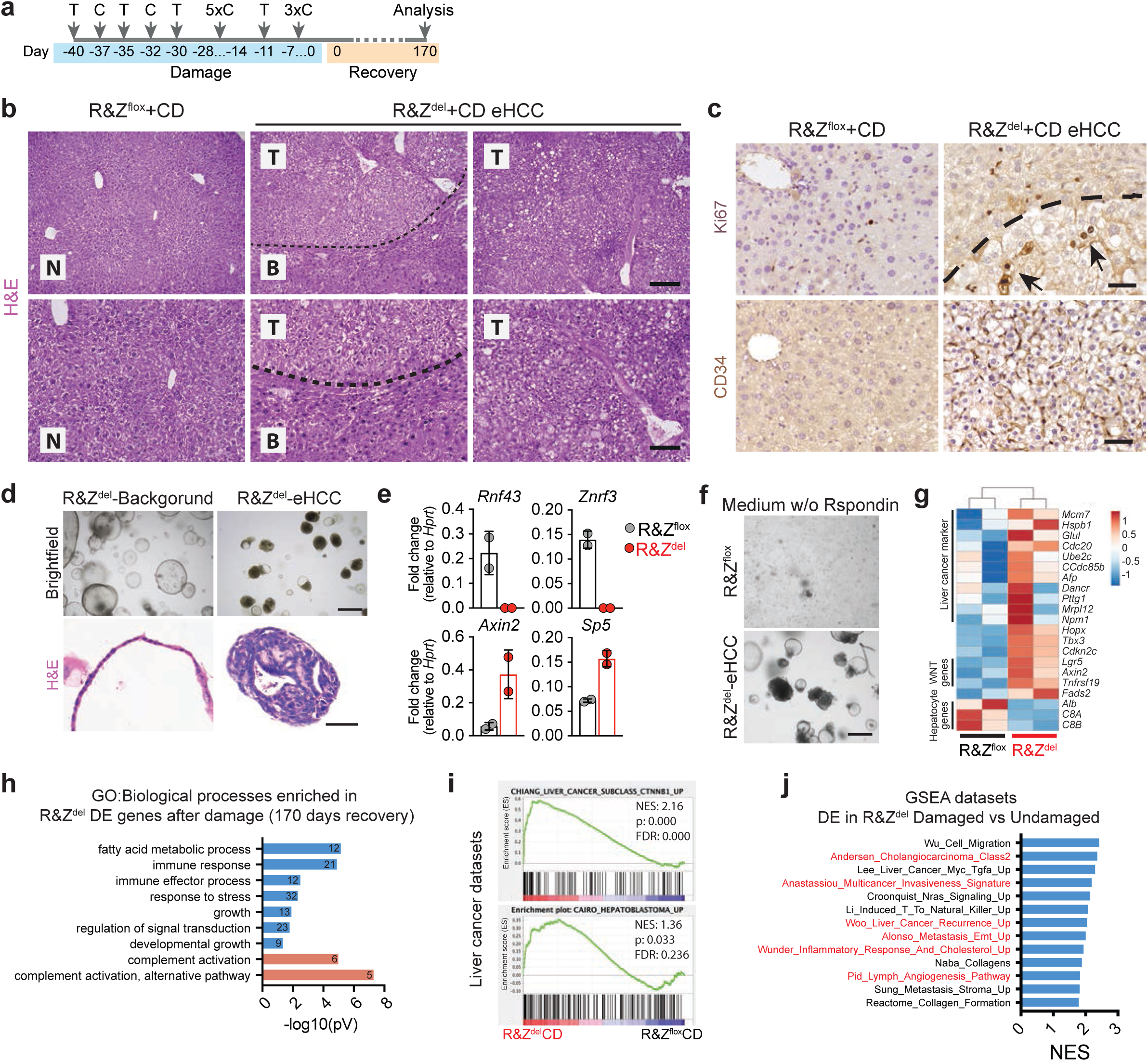
Liver-specific *Rnf43 and Znrf3* (R&Z) deletion leads to formation of early hepatocellular carcinoma after CCl4-induced chronic damage. Liver-specific R&Z mutant mice were injected with CCl4 for 6 weeks to induce chronic damage to the liver and tissues were collected 170 days later. Histopathological analysis revealed the presence of several lesion characterized as early hepatocellular carcinoma in 66.7% of the mice analyzed. R&Z^flox^ mice lacking the AlbCreERT2 allele were also injected with tamoxifen and CCl4 and used as controls. **a)** Timeline of the experiment. T, tamoxifen; C, CCl4. **b-c)** Histopathological analysis revealed presence of early hepatocellular carcinomas. Dotted black lines mark eHCC border. **b)** Representative pictures of H&E stainings of tumours and background tissues. N, normal tissue; B, background tissue; T, tumoural tissue. Scale bar, 200μm (H&E top panel); 100μm (H&E bottom panel). **c)** Tumour presented increased proliferation (Ki67) and vascularisation (CD34). Arrows, Ki67^+^ cells. Scale bar, 50μm. **d)** Brightfield pictures and H&E staining of tumouroids isolated from an eHCC lesion. Scale bar, 500um for brightfield pictures and 50 μm for H&E. **e)** qPCR expression analysis for WNT target genes in organoids from R&Z^flox^ and R&Z^flox^ eHCC. Data represent mean +/- SD of n=2 experiments. **f)** Brightfield pictures of organoids cultured without Rspondin. Scale bar, 200 μm. **g)** Heat-map analysis of the RPKM values (raw z-scored) of liver cancer markers, WNT target genes, hepatocyte differentiation genes and others. **h)** Graph showing selected gene ontology (GO) terms significantly enriched for genes up-(blue) and down-(pink) regulated in R&Z^del^ compared to R&Z^flox^. The numbers denote the number of genes associated to each term. Full list of significant terms can be found in **Supplementary Data Sets 1. i)** GSEA revealed that 14 out of 28 gene sets involved in liver cancer and NASH were significantly enriched in our DE gene signatures (FDR<25%). Representative plots for enriched sets are shown. Details of all the positively and negatively correlated signatures are given in **Supplementary Data Sets 1. j)** Graph showing normal enrichment score (NES) for selected GSEA datasets significantly enriched (FDR<25%; p<0.05) in R&Z^del^ damaged livers against R&Z^del^ undamaged livers. Full list can be found in **Supplementary Data Sets 1**.

To specifically better characterize the HCC lesions, we microdissected the tumour and either processed them for molecular analysis or cultured them in our previously described human primary liver cancer organoid method, that allows the *in vitro* expansion of HCC tumour cells derived from well/moderate differentiated HCC, but not from very well differentiated HCC tissue^42^. Molecular analysis of the lesions indicated that the HCC tumours were indeed devoid of *Rnf43/Znfr3* expression and expressed high levels of *Axin2*, confirming the *Rnf43/Znrf3*^*del*^ origin of the tumour lesion (**Supplementary Fig. 6d**). Interestingly, HCC lesions but not the background liver, expanded as tumour organoids *in vitro* by forming compact structures composed of tumoural cells that filled in the lumen of the organoid and expressed high levels of WNT target genes (*Axin2* and *Sp5*) (**Fig. 5d-e**). Of note, *Rnf43/Znrf3*^*del*^ tumour organoids, but not WT controls, grew long-term (up to passage 10) in the absence of Rspo1 ligand in the medium, confirming their origin from *Rnf43/Znrf3*^*del*^ cells (**Fig. 5f**).

To gain detailed insight on the molecular signature of these tumours, we performed RNAseq analysis of the whole tissue and compared it to WT littermate controls subjected to the same regimen and recovery time. We identified 214 differentially expressed genes (**Supplementary Fig. 3a-b**). Amongst the upregulated genes we found stem cell (*Hopx*^43^, *Tnfsrf19*^44^) and early hepatoblast (*Tbx3*^14,45^ and *Lgr5*^46^) markers, as well as genes involved in lipid metabolism, such as *Fads*. Conversely, mature hepatocyte markers like *C8A* and *C8B* and *Alb*, were significantly downregulated (**Fig. 5g** and **Supplementary Data Set 1**). Interestingly, we also found upregulation of several well-known liver cancer markers including *Mcm7, Cdc20, Afp* and *Hspb1* among others, in agreement with the neoplastic transformation of the tissue (**Fig. 5g**). Similarly, GO and GSEA analysis revealed enrichment on fatty acid metabolism (**Fig. 5h**), and several cancer signatures, including tumours with β-catenin activation^47^, hepatoblastoma^48^ and cancer patients with NASH (**Fig. 5i** and **Supplementary Fig. 6e**)^29^. Interestingly, when comparing the *Rnf43/Znrf3*^*del*^ livers exposed to CCl4 to undamaged *Rnf43/Znrf3*^*del*^ controls, we found significant enrichment with advanced cancer and metastatic cancer datasets, as well as with inflammation and cholesterol uptake (**Fig. 5j**). Similar enrichment was also detected at an earlier time point (at 50 days recovery) (**Supplementary Fig. 6f**). These results indicated that the progression to malignancy from a steatotic/NASH state requires the activation of a damage-regenerative response in *Rnf43/Znrf3*^*del*^ mice.

Altogether, this data indicated that RNF43/ZNRF3 are not primary tumour suppressors in liver, yet their deletion predisposes to cancer due to altered lipid metabolism and improper tissue regeneration upon chronic damage.

### Hepatocellular carcinoma patients bearing *RNF43/ZNRF3* mutations exhibit altered lipid metabolism and poorer prognosis

To determine the relevance of our findings in human liver cancer, we next sought to assess the effect of *RNF43/ZNRF3* mutations in human primary liver tumours. For that, we took advantage of the publicly available ICGC data collection (project LIRI-JP)^49,50^, where both genomic and expression data of the same tumours is accessible (**Fig. 6a**). LIRI-JP harbours 26 HCCs with mutations in *ZNRF3* and no other mutations in the WNT pathway (ZNRF3 tumours) and 64 HCCs with wild-type copies of the gene (WT tumours). We found only 9 tumours with *RNF43* mutations. Therefore, due to the low number on RNF43 mutant patients, for the following analysis, we focused on HCCs with *ZNRF3* mutations.

**Figure 6.**
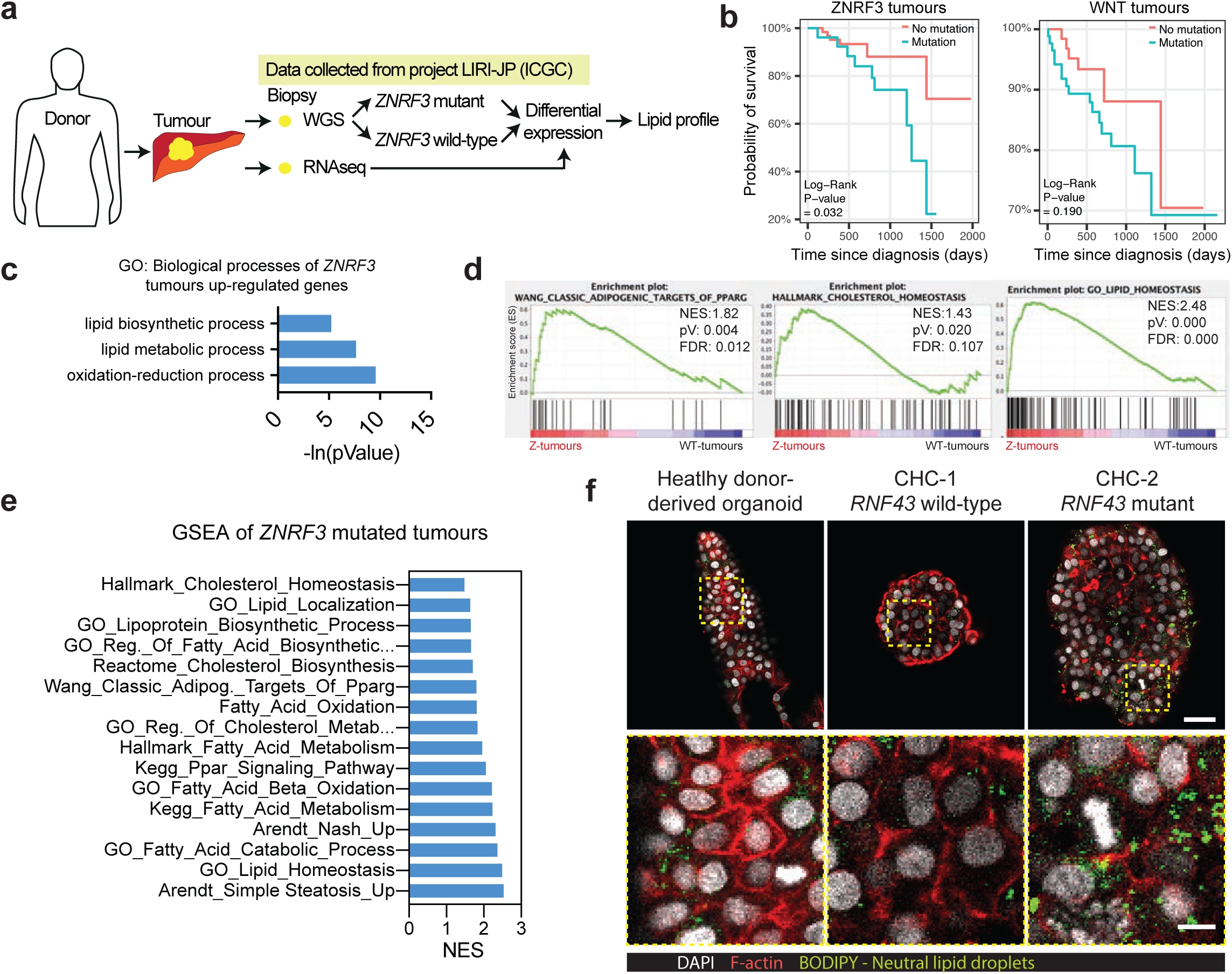
Human hepatocellular carcinomas mutated in ZNRF3 present lipid metabolic alterations and poor prognosis. **a-e)** Clinical data, whole genomic sequencing and RNAseq data were downloaded from ICGC database (project code LIRI-JP) and used to determine the prognosis as well as to compare the expression pattern of human tumours mutated in ZNRF3. **a)** Schematic representation of experimental procedures. **b)** Graph showing survival curve for HCC patients with ZNRF3 mutations or with mutations in the WNT pathway. **c)** Graph showing some of the GO terms significantly enriched for genes up-regulated in ZNRF3-mutated human tumours. Full list is shown in **Supplementary Data Sets 4. d-e)** GSEA revealed that 81 out of 207 gene sets involved in lipid metabolism were significantly enriched in our DE gene signatures (FDR<25%). Representative plots for lipid metabolism enriched sets are shown **(d)**. Details of all the positively and negatively correlated signatures are given in **Supplementary Data Sets 4. f)** Human liver organoids derived from a healthy donor or from patients with liver cancer (mixed subtype, CHC) were grown in our standard tumoroid medium and collected and processed for neutral lipids staining (Bodipy, Green) as described in methods. CHC-1 is WT while CHC-2 is mutated for RNF43 (Leu418Met). Note that there is an increase in the number of intracellular lipid droplets in the mutant CHC2 compared to CHC1 or donor. Red, actin staining. White, Dapi. Scale bar, 60μm (top panel); 15μm (bottom panel).

Interestingly, we observed that patients bearing ZNRF3 tumours had significantly poorer prognosis than patients with WT tumours (**Fig. 6b, left panel**). Using multivariate survival analysis, we showed this result was independent of mutational burden, clinical tumour stage and alterations in the WNT pathway (*β-catenin, APC, AXIN1*) (WNT tumours) (**Supplementary Table 1**). No significant decrease in survival rates was observed for patients with mutations in other components of the WNT pathway, consistent with what had been previously reported^22^ (**Fig. 6b, right panel**). Similarly, no cumulative effect of ZNRF3 tumours over WNT tumours was observed (**Supplementary Table 1**).

To further explore the implication of *ZNRF3* mutations in human liver cancer, we took advantage that for 18 of the ZNRF3 tumours and for 48 of the WT tumours we could also obtain associated transcriptomic data. Hence, we then performed differential expression analysis of the livers of patients harbouring *ZNRF3* mutations compared to liver cancer patients with the *ZNRF3* WT allele. We found 104 genes up regulated and 1073 genes downregulated. Notably, amongst the upregulated genes we found genes involved in fatty acid (*FASN, AKR1B10*), Fatty-Acyl-CoA (*THEM5)* and cholesterol (*TM7SF2)* biosynthesis, activators of lipogenesis (*MLXIPL)* as well as genes involved in intracellular lipid storage *(PLIN4, CIDEC, CD36-like 1*) (**Supplementary Data Sets 4**). In line with this, GO analysis revealed enrichment of terms involved in lipid metabolism, such as ‘lipid biosynthetic process’ and ‘lipid metabolic process’ (**Fig. 6c**) while GSEA confirmed a positive enrichment for datasets involved in lipid metabolism such as ‘Adipogenic targets of PPARG’, ‘cholesterol homeostasis’ and ‘lipid homeostasis’ (**Fig. 6d-e**). Of note, some of these datasets were also enriched in our *Rnf43/Znrf3*^*del*^ mice (see **Fig. 2e** and **Fig. 6d-e**).

To determine whether the patient’s lipid phenotype would be the result of lipid accumulation in a cell-autonomous manner, we took advantage of our tumour-organoid culture system that enables the *in vitro* expansion of primary tumour cells from both HCC and mixed subtype (CHC)^42^. None of our HCC organoid lines harboured *ZNRF3* mutations. However, we found that our CHC-2 tumour organoid line presented an homozygous (deleterious) *RNF43* mutation (Leu418Met). Notably, this CHC-2 mutant line accumulated intracellular lipid droplets even in the absence of any fatty acid supplementation in the medium (**Fig. 6f**, right panel). Contrary, tumour organoid derived from either liver cancer patients without mutations in any of the two genes (CHC-1) or donor-derived organoids (derived from non-cancer tissue) presented less intracellular lipid droplets (**Fig. 6f**, left panel and middle panel).

In summary, our results indicate that human liver cancer patients bearing *ZNRF3* mutations present poorer prognosis and a significantly altered liver lipid metabolism, the latter in part due to the accumulation of lipid droplets in the tumour cells.

## Discussion

In the liver, several members of the Wnt pathway had been reported recurrently mutated in hepatocellular carcinoma (HCC) (32.8% beta-catenin, 15.2% *AXIN1* and 1.6% *APC*)^51^. Recently, the E3 ligases RNF43 and ZNRF3 were added to the list of liver cancer mutated genes^22,23^, however, whether they act as *bona-fide* tumour suppressor genes, passenger mutations or predisposing risk factors remained unexplored. Here we describe that in the liver, *Rnf43/Znrf3* do not drive, yet predispose, to cancer by altering the lipid metabolic rheostat and impairing the capacity of hepatocytes to regenerate upon chronic damage. Our results indicate that *Rnf43/Znrf3* are a subtype of tumour suppressor genes that when mutated lead to a significant susceptibility to cancer, resembling landscaper genes, which contribute to neoplastic transformation by altering the tissue environment^52^. We found that deletion of both homologues in mouse results in altered lipid metabolism, steatotic foci and hepatocyte injury, thus resembling human NASH. Remarkably, all these metabolic changes were cell-autonomous and, hence, present in mice fed normal diet, without the need of exogenous fatty acid supplementation, contrary to what is described for most NASH mouse models^53^. Upon chronic damage, *Rnf43/Znrf3* mutant livers presented a defect in the maturation of the *de novo* formed hepatocytes, which resulted in extensive fibrosis, a cirrhotic phenotype and subsequently led to the development of HCC. Remarkably, our results translated to liver cancer patients bearing ZNRF3/RNF43 mutations as we found that these sub-groups of patients present cell-autonomous lipid accumulation in the tumour cells and poorer survival compared to patients with wild-type ZNRF3/RNF43 alleles.

Our findings might seem at odds with reported results whereby WNT3a injection prevents lipid accumulation in mice with active non-canonical WNT through an LRP6 receptor mutation (LRP6^R611C^)^54,55^ and with the phenotype of the core canonical WNT component APC in the liver, which results in HCC without the need of any additional hit or tissue damage^37^. However, recent reports have demonstrated a link between WNT signal strength and pathway outcome, suggesting that the strength of activation of the pathway as well as the nuclear beta-catenin fold change, rather than absolute levels, are critical for target gene activation^56^, stem cell behaviour or malignant transformation^57–59^. In that regard, RNF43/ZNRF3 mutant colorectal cancer tumours have been found to strictly depend on exogenous WNTs^21^, unlike *Apc, Axin1/2*, or *βcatenin* mutants that are WNT independent. Similarly, it has been recently shown that *Rnf43/Znrf3* malignant cells are still sensitive to small molecule inhibitors that would act at the receptor/ligand level and prevent cancer progression^21,60,61^. In that regard, our data whereby porcupine inhibition facilitates the hepatic differentiation of *Rnf43/Znrf3* mutant organoids suggests that inhibiting WNT production could facilitate the *in vivo* resolution of the otherwise impaired regenerative process in the *Rnf43/Znrf3* mutants undergoing tissue damage. This leads us to speculate that for the development of liver cancer in RNF43/ZNRF3 mutants, the activation of Wnt secretion during the damage-response might be a crucial step towards the malignant progression. Since a number of components of the Wnt pathway are mutated in liver cancer, identifying those individuals that could benefit from treatments with WNT inhibitors could significantly influence the therapeutic intervention and management of liver cancer patients.

On another note, we found that the altered lipid accumulation was dependant on canonical Wnt and cell-autonomous due, in part, to increased intracellular lipid uptake. CD36, as well as high fat intake, have been recently described in oral cancer as essential factors that determine the probability of a cancer cell to become metastatic^62^. In addition, CD36 was also a direct canonical Wnt target (see **Fig. 2**). Hence, it is tempting to speculate that those liver cancer patients bearing RNF43/ZNRF3 or any other canonical WNT mutation are at higher risk of developing metastatic cancer than patients with the corresponding wild-type alleles. In that regard, we found that ZNRF3 mutant liver cancer patients present decreased survival rates and lipid metabolic alterations, including overexpression of the CD36-like 1 (*SCARB1*) receptor.

In summary, our results provide a framework for understanding the role of RNF43/ZNRF3 as predisposing risk factors for the development of liver cancer, whereby the increase in intracellular fat and impaired regenerative capacity result in cirrhosis that progresses towards malignancy upon tissue damage. With the alarming increase in the consumption of saturated fats and sugar world-wide, recognizing those individuals with mutations in these E3 ligases or any other canonical WNT signaling component might facilitate the discovery of populations at risk of developing liver steatosis, NASH, cirrhosis or cancer^63^. In that regard, our results indicate that while RNF43/ZNRF3 mutant organoids already and cell-autonomously accumulate lipid, this uptake is increased upon free-fatty acid supplementation, calling for the intervention on reducing lipid/fat intake as a preventive therapeutic strategy for those RNF43/ZNRF3 mutant individuals at risk of developing liver disease. Identifying these subgroups at risk would have strong implications for the clinical management of liver disease, liver cancer patients, patients with steatosis or even healthy individuals with mutations in either of the two homologues or in other Wnt signaling components, who could be advised low fat diet and/or more frequently monitored for liver plasma markers.

## Methods

### Animals

All mouse experiments were conducted in accordance with procedures approved by the UK Home Office relating to the use of Animals in research (Animals Act 1986). Conditional knock-out mice for *Rnf43* and *Znrf3* (*Rnf43/Znrf3*^*flox*^) were previously generated and described in Koo et al. 2012^18^. Briefly, exons encoding for the ring domain of *RNF43* and *ZNRF3* were flanked by LoxP sites to drive excision in presence of a CRE protein (**Supplementary Fig. 1c**). *Rnf43/Znrf3*^*flox*^ mice were crossed with either Alb1-CreERT2^24^ or Sox9-CreERT2 (Jackson Laboratory JAX - http://jaxmice.jax.org/) to generate Alb1-CreERT2-R&Z^flox^ (used for the *in vivo* studies) and Sox9-CreERT2-R&Z^flox^ (used to generated R&Z^del^ organoids). The Cre enzyme was induced by intraperitoneal injection of 200mg/kg tamoxifen (Sigma-Aldrich) dissolved in sunflower oil. Sunflower oil only was injected to control mice. Alternatively, deletion was performed after tail vein injections of 2.5 × 10^11^ vg/ml of AAV.TBG.PI.Cre.rBG. Control mice received pAAV.TBG.PI.Null.bGH. AAV8 viruses were diluted in sterile PBS to a final volume of 100ul. AAV.TBG.PI.Cre.rBG and pAAV.TBG.PI.Null.bGH were a gift from James M. Wilson (Addgene plasmid # 107787; http://n2t.net/addgene:107787;RRID:Addgene_107787 and Addgene plasmid # 105536; http://n2t.net/addgene:105536;RRID:Addgene_105536). Liver damage experiments were performed using CCl4 (Sigma-Aldrich) diluted in corn oil and administrated 500ul/kg (CCl4 dilution 1:4) multiple times according to protocol (**Fig. 3a** and **Fig. 5a**). Mice were sacrificed by either cervical dislocation or carbon dioxide overdose and blood was collected by cardiac puncture or tail vein prick.

For hepatocyte collection and cell sorting, mice were culled by carbon dioxide intoxication and head was removed to allow exsanguination. Livers were perfused with 10ml of perfusion buffer (0.5mM EGTA in PBS), then with 10ml of perfusion buffer with BSA (Sigma-Aldrich) and then with 10ml of pre-warmed 0.5mg/ml Collagenase A (Sigma-Aldrich) solution (HBSS 1x, 5mM CaCl2, 20mM HEPES). After 2 minutes incubation, livers were carefully removed, place in STOP solution (HBSS 1x, 5mM CaCl2, 20mM HEPES, 0.5% BSA) and disaggregated using tweezers. Mixture was washed several times in STOP solution (50G for 2 minutes) and filtered with a 100μm strainer. Pellet containing hepatocytes was resuspended in RLT lysis buffer (Quiagen) and frozen at −80C for RNA extraction. Supernatant was centrifuged again at 300g for 5 minutes to collect non-parenchymal cells. Pellet was divided in two fractions and stained either with EpCAM-APC (eBioscience) and CD31-PE-Cy7 (AbCAM) antibodies or EpCam-APC and CD11b-PE-Cy7 (BD Bioscience) antibodies. Positive cells were sorted using a Flow Sorter MoFlo (Dako Colorado, Inc.), resuspended in RLT lysis buffer and stored at −80C.

### Histology

Tissues were dissected and then fixed overnight in 10% buffered formalin (Sigma-Aldrich). For paraffin embedding, tissues were dehydrated in increasing concentration of ethanol (70%, 95% and 100% for 2h each) ending in xylene and then incubated overnight in paraffin (Sigma-Aldrich). The following day, tissues were embedded in paraffin blocks and subsequently sectioned at a thickness of 5μm using a microtome. Sections were mounted onto SuperFrost Plus slides (ThermoFisher) and incubated from 3 to 12h at 60C before staining. H&E (Sigma-Aldrich), Oil Red O (Sigma-Aldrich) and PicroSirius Red (AbCAM) staining were performed according to manufacturer instruction. For immunoistochemistry, sections were deparaffinised in two changes of xylene 5 minutes each and rehydrated in descending grades of ethanol (100% 2x, 95%, 70%) 3-5 minutes each ending in water. Antigen retrieval was performed either enzymatically or heat-mediated based on primary antibody (see **Supplementary Data Sets 5**). For enzymatic retrieval, sections were incubated 10 minutes at 37C and 10 minutes at RT with TE buffer containing 0.5% Triton X-100 and 1/1000 proteinase K (800U/ml). For heat-mediated antigen retrieval, samples were incubated in either citrate buffer (10mM Sodium Citrate, pH 6) or Tris-EDTA buffer (10mM Tris base, 1mM EDTA solution, 0.05% Tween 20, pH 9.0), heated in a microwave to 95-98C for 20 minutes and then cooled at RT for 20 minutes. Afterwards, samples were blocked and permeabilized for 1.5 hours using TBS buffer containing 2% foetal bovine serum (FBS), 1% bovine serum albumin (BSA) and 1% or 0.1% of Triton x-100 based on antibody (see Table 7.2 for details). Samples were than incubated overnight with primary antibody at antibody-dependent concentration (see **Supplementary Data Sets 5**). The following day, samples were washed in TBS and endogenous peroxidases were inactivating incubating 15 minutes in methanol containing 3% of hydrogen peroxide. Few drops of Poly-HRP-GAMs/Rb IgG (Immunologic) secondary antibody was added directly on the sections and incubated for 30 minutes. After washing, signal was developed using Bright Dab substrate kit (Immunologic) using manufacturer instruction. Sections were counterstained in hematoxylin for 5 minutes, dehydrated in ethanol (70%, 95% and 100% 2x) and in two changes of xylene 5 minutes each and finally mounted using xylene-based DPX mounting media (ThermoFisher). Slides were analysed using optical microscopy. Quantification of staining for GS, Sirius Red and PCK were performed by measuring stained area using ImageJ (Fiji).

### RNA analysis

RNA was extracted from tissues, cells or organoid cultures using the RNeasy Mini RNA Extraction Kit (QIAGEN) according to the manufacturer’s protocol and reverse transcribed with the Moloney Murine Leukemia Virus reverse transcriptase (M-MLVRT) (Promega) as follows: 10 minutes at room temperature, 50 minutes at 50C and 15 minutes at 70C. The resulting cDNA was amplified with the iTaq™ Universal SYBR. Green Supermix (Bio-Rad) and a desired primer pair on the CFX Connect™ Real-Time PCR Detection System (Bio-Rad). The cycling conditions used were: 95C for 3 minutes and 39 cycles of 95C for 10 seconds and 55-60C for 30 seconds (depending on the primer pair), followed by a melting curve (from 55-60C to 95C with an increase of 0.5C per cycle) to ensure amplicon specificity. Gene expression was normalised to the expression of the housekeeping gene Hprt. Primers used are listed in **Supplementary Data Sets 5**.

### RNA sequencing

RNA libraries were prepared for sequencing using the Smart-Seq2 protocol^64^. RNA-Seq libraries were sequenced on an Illumina HiSeq 4000 instrument in single read mode at 50 base length. Reads were filtered for low quality (<Q20) with Sickle (version 1.33)^65^. Reads were then mapped to mm10 UCSC reference genome^66^ using the STAR aligner (version 2.5.0a)^67^ with the parameters “--outSAMmultNmax 1, --quantMode GeneCounts”. Raw counts were generated using featureCounts (version 1.6.0)^68^ software and includes all exons for a gene from the mm10 UCSC GTF file.

### TCF4 Binding analysis

Location of TCF4 peaks were taken from a previous study^16^ and annotated using ChIPseeker (version 1.12.1)^69^ in R on the mm9 genome. Genes were filtered to include those with a peak within 5kb up or downstream of their transcriptional start site. We defined a lipid metabolism set of genes by combining genes contained in the GSEA hallmark fatty acid metabolism gene set, the Reactome cholesterol biosynthesis gene set, Wang classic adipogenic targets of Pparg and the gene ontology lipid metabolic process (including child terms) set. Random peak sets were generated by shuffling the peak locations using ChIPseeker. This was done 1,000 times to generate empirical p-values for the enrichment of lipid metabolism genes with TCF4 binding sites. Venn diagrams were generated using the web tool Venny 2.1 (http://bioinfogp.cnb.csic.es/tools/venny/).

### RNA-seq differential expression analysis

For each comparison expressed genes were defined as having counts per million of at least 1 in half of the samples. Normalisation was done using calcNormFactors and variance estimators calculated using estimateDisp in edgeR. Differential expression analysis was performed using edgeR (version 3.18.1)^70^ and significantly differential expressed genes were defined as having a false discovery rate (FDR) less than 10% and an absolute log2 fold change greater than 1. Empirical p-values for enrichment of TCF4 regulated genes in those showing differential expression was similarly calculated using the random peak sets used in the TCF4 peak analysis.

### Heatmaps, gene ontology and GSEA

Heat map analysis were performed by submitting RPKM values to web tool ClustVis (https://biit.cs.ut.ee/clustvis/). GSEA was performed using the GSEA 3.0 software downloaded from http://software.broadinstitute.org/gsea/index.jsp. GO analysis was performed using webtool DAVID (https://david.ncifcrf.gov).

### Organoid cultures

Liver organoids were derived and cultured as previously described^32^. *Rnf43/Znrf3*^*del*^ organoids were generated by isolating cells from *Sox9CreERT-Rnf43/Znrf3*^*flox*^ mice and then treated *in vitro* with 3μM 4-hydroxy-tamoxifen (Sigma) to induce deletion. For the isolation, livers were collected in PBS and dissociated using a collagenase solution (Collagenase type XI 0.012%, dispase 0.012%, FBS 1% in DMEM medium) and incubated for 3 to 4 hours at 37°C. Mixture was washed several times and ducts were hand-picked, mixed with Matrigel (BD Bioscience) and seeded in a 24 multi-well plate. After Matrigel had polymerized, culture medium was added. Culture conditions for expansion (EM) were based on AdDMEM/F12 (ThermoFisher) supplemented with 1% B27 (Invitrogen), 1% N2 (Invitrogen), 1.25mM N-acetylcysteine (Sigma-Aldrich), 10nM gastrin (Sigma-Aldrich), 50ng/ml mEGF (Peprotech), 5% RSPO1 conditioned medium (homemade), 100ng/ml Fgf10 (Peprotech), 10mM nicotinamide (Sigma-Aldrich) and 50ng/ml HGF (Peprotech). To establish the culture, media was supplemented with 25ng/ml Noggin (Peprotech), 30% WNT conditioned medium (home-made) and 10μm Rho-kinase inhibitor (Y27632, Sigma-Aldrich), for the first week. To generate *Rnf43/Znrf3*^*del*^ tumouroids from eHCC lesions, RSPO1 and WNT conditioned medium were removed from culture conditions. Weekly, organoids were removed from the matrigel, mechanically dissociated with a narrowed Pasteur pipette, and transferred to fresh matrix in a 1:4 split ratio.

Human tumouroids were cultured as previously described^42^ in AdDMEM/F12 (ThermoFisher) supplemented with 1% B27 (Invitrogen), 1% N2 (Invitrogen), 1.25mM N-acetylcysteine (Sigma-Aldrich), 10nM gastrin (Sigma-Aldrich), 50ng/ml mEGF (Peprotech), 10% RSPO1 conditioned medium (home-made), 100ng/ml Fgf10 (Peprotech), 10mM nicotinamide (Sigma-Aldrich), 25ng/ml HGF (Peprotech), 10 μM forskolin and 5 μM A8301.

### Differentiation and staining

To differentiate cultures into hepatocyte-like cells, liver organoids were cultured and differentiated as previously described^32^. Organoids were collected, disrupted by pipetting, seeded in matrigel and cultured in complete expansion medium for four days. On day 5, medium was changed to differentiation medium (DM) based on AdDMEM/F12 (ThermoFisher) supplemented with 1% B27 (Invitrogen), 1% N2 (Invitrogen), 1.25mM N-acetylcysteine (Sigma-Aldrich), 50ng/ml mEGF (Peprotech), 50nM A8301 (Tocris), 10uM DAPT (Sigma) and 100ng/ml Fgf10 (Peprotech). Medium was refreshed every two days. Organoids were cultured in DM for a total of 9 days. Dexamethasone (3uM) was added to the differentiation medium at the end of the protocol for a total of three days. To enhance differentiation of *Rnf43/Znrf3*^*del*^ organoids, DM medium was supplemented with IWP2 (3uM) and iCRT3 (25 μM) to make DMWi medium.

For lipid treatment, medium was supplemented with 500uM of oleic acid and it was replaced every two days for a total of 4 days. Organoids were collected using a plastic pasteur pipette, washed with cold PBS several times to remove matrigel or BME carefully to not disrupt their 3D structure. Subsequently, organoids were fixed for 40 minutes on ice with 10% Formalin. Fixed organoids were stored at 4°C. For staining, fixed organoids were blocked with 1% Bovine Serum Albumin (BSA), 2% Donkey serum (DS), 1% DMSO in PBS for 2h at RT. Primary antibody was incubated o/n at 4°C in 0.05% BSA in PBS. The secondary antibodies were added at a concentration of 1:150 for 2h at RT. Nuclei were stained with Hoechst 33342 (Invitrogen) at a dilution 1:1000 for 15min at RT. To assess lipid accumulation, cells were stained using BODIPY 493/503 dye (2uM in PBS). When combined with a membrane staining, BODIPY was added after the incubation with the secondary antibody. BODIPY was incubated for 3h at RT protected from light. Imaging was performed using an SP8 White Light inverted confocal microscope. All images were acquired using a 63x water immersion lens. The images were edited and analyzed using the Leica Application Suite X software (LAS X, v1.5.1.1387) and ImageJ (v1.51j8).

### ZNRF3 Human mutation analysis

We downloaded RNA-seq data for the liver cancer hepatocellular carcinoma study from Japan (LIRI-JP) from the ICGC database^49,50^. The effect of *ZNRF3* mutation on patient survival was assessed using survival analysis. For the LIRI-JP data set the survival data was downloaded from ICGC. There were 260 donors with clinical data in the LIRI-JP study. Mutation data was also downloaded from ICGC and annotated using SNP nexus^71^. Mutations were filtered based on their predicted consequence type under different metrics from SNP nexus. A mutation was included if it met the criteria for any one of the following metrics: 1) SIFT prediction was classed as damaging with confidence “high”; 2) Polyphen predicition was either possibly or probably damaging; 3) Eigen PC score was greater than zero; 4) FatHMM non-coding score was positive; 5) Fit conservation score was greater than 0.2. Patients harbouring mutations in either *APC, AXIN1* or *CTNNB1* were considered to have a mutation in WNT, regardless of their *ZNRF3* mutation status. Wild-type (WT) patients were defined as having neither a WNT, *ZNRF3* or *RNF43* mutation (n=64). Patients were classed as having *ZNRF3* mutation only when the *ZNRF3* mutation occurred without the presence of WNT (n=26). We then tested for differences in survival between patients with *ZNRF3* mutation only and wild-type patients. Differences between the survival curves were tested using the log-rank test and Kaplan-Meier curves plotted using the survival (version 2.41-3) and GGally (version 1.3.2) packages in R. In the multiple survival regression, we included the ZNRF3 mutation status, RNF43 mutation status as well as WNT mutation status and an interaction term between ZNRF3 and WNT mutation. To include tumour stage in the survival analysis we used the tumour stage at diagnosis information in the clinical data. Patients were split into two groups, those whose tumour stage was either 1 or 2 and those whose tumour stage was 3 or 4. We also included sex as a predictor. Mutational burden was estimated from the mutation data downloaded from the ICGC database as number of mutations, excluding those in intergenic regions. Analysis was performed in R using the survival package and the cox proportional hazards regression model. The LIRI-JP dataset contains RNA-seq data with matching normal solid tissue and tumour samples for a set of 196 donors. There are 402 samples in total, each donor has at least one tumour and one normal control sample. Genes were filtered to remove those not expressed where expressed genes are defined as having counts per million greater than 1 in more than 80 samples. These samples were then stratified according to their mutation status. As with the survival analysis we modelled differences in gene expression between wild-type and ZNRF3 only patients using a multiple regression model in R. There were 48 donors with wild-type mutation status and matching tumour and normal RNA-seq samples and 18 donors classed as having *ZNRF3* mutation only with matching tumour and normal RNA-seq samples. The regression model contained three terms, one for specimen type (tumour or normal), one for *ZNRF3* mutation status and the third for donor ID that accounted for patient variability^70^. Normalisation of RNA-seq samples was done using calcNormFactors and variance estimators calculated using estimateDisp in edgeR. Genes were defined as differentially expressed between the tumour samples of wild-type patients compared to those with *ZNRF3* mutation only at logFC>+/-1 and 10% FDR following multiple hypothesis correction^72^.

## Supporting information

Supplementary Dataset 1 - RNAseq

Supplementary Dataset 2 - Lipid analysis

Supplementary Dataset 3 - Serum analysis

Supplementary Dataset 4 - Genetic human analysis

Supplementary Dataset 5 - List of materials

## Data availability

All RNAseq data are available at the Gene Expression Omnibus (GEO) under accession number GSE133213. Token number: kjmvyeeylbqzfil.

## Acknowledgements

We would like to acknowledge the Gurdon Institute and Cambridge Stem Cell Institute Animal facilities for their support. The Gurdon Institute Imaging Facility for microscopy and image analysis support. Dr Andy Riddell (Cambridge Stem Cell Institute) for cell sorting and Dr Maike Paramor (Cambridge Stem Cell Institute) and Mr Kay Harnish of the Gurdon Institute genomics facility for assistance with library preparation and high-throughput sequencing, respectively. Keerthi Sindhura for initial bioinformatics analysis. Dr Laura Broutier for help on the early phases of the project; Dr Luigi Aloia for WNT target genes expression analysis after damage and Lucia Cordero Espinoza for the preparation of sorting experiment. M.H. is a Wellcome Trust Sir Henry Dale Fellow and is jointly funded by the Wellcome Trust and the Royal Society (104151/Z/14/Z); M.H. and G.M. are funded by a Horizon 2020 grant (LSFM4LIFE). In the early phase of the project, G.M. and B.K.K. were funded by a Marie Curie Initial Training Network (Marie Curie ITN WntsApp 608180). The authors acknowledge core funding to the Gurdon Institute from the Wellcome Trust (092096) and CRUK (C6946/A14492).

## Author Contributions

M.H. and B-K.K. conceived and G.M., B-K.K and M.H. designed the project. G.M., S.K. and M.H. designed and performed experiments and interpreted results. G.M. prepared and analysed RNAseq and C.P. and C.R.B. performed and interpreted all bioinformatic analyses. R.A-B. and G.M performed and S.D. analysed all the histological sections. G.M. and M.H. wrote the manuscript. All authors read and commented on the manuscript.

---

**Supplementary figure 1.**
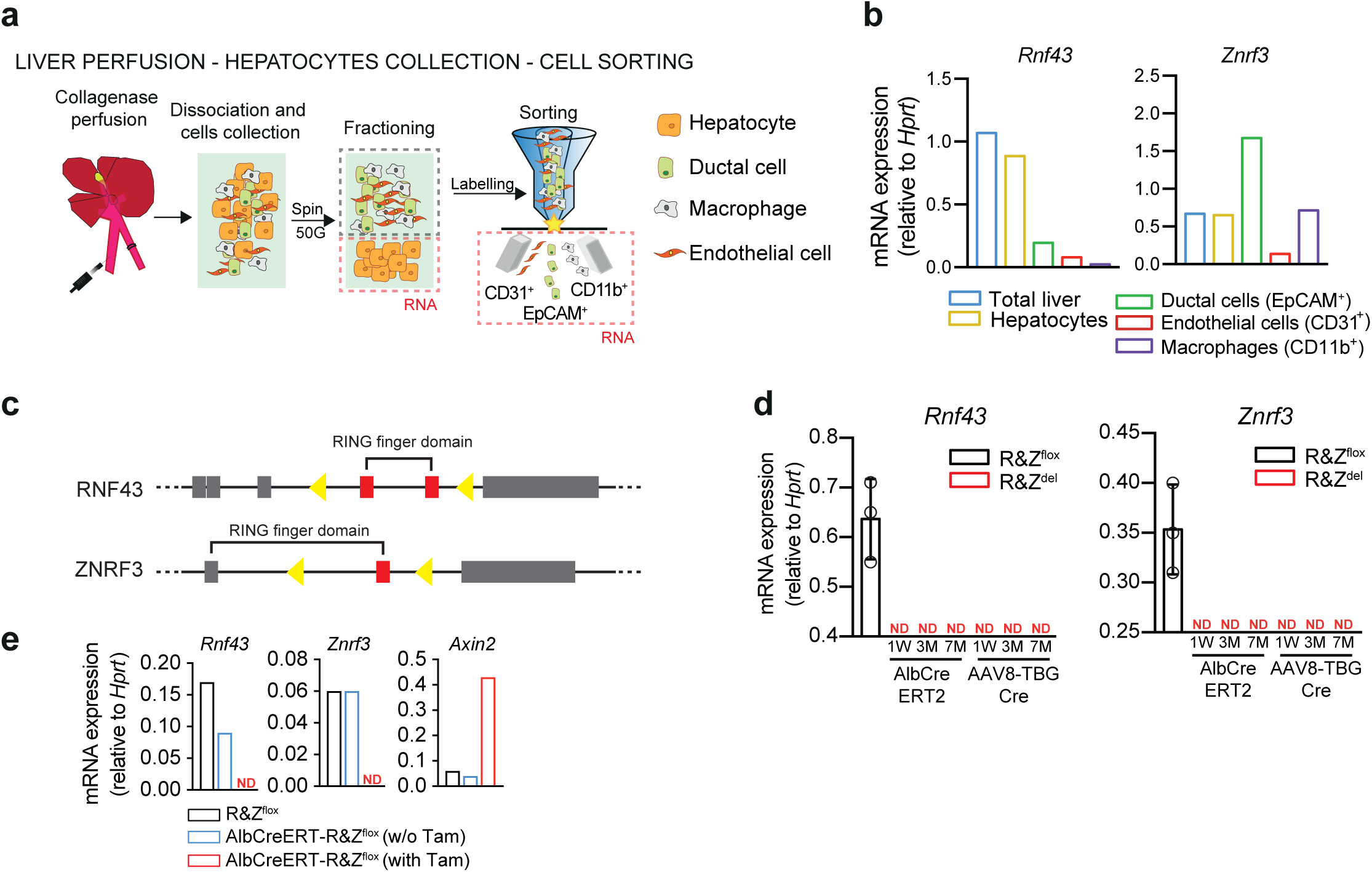
*Rnf43 and Znrf3* (R&Z) are expressed in the liver and can be efficiently deleted after tamoxifen or AAv8-TBG-Cre injection. **a)** Diagram representing strategy for liver perfusion, hepatocyte collection and cell sorting. Livers were perfused with collagenase and then mechanically dissociated. Hepatocytes were separated from the mixture by centrifugation. Ductal cells were labeled with anti-EpCAM antibody, macrophages with anti-CD11b antibody and endothelial cells with anti-CD31 antibody and then sorted. RNA was collected from all the isolated cell types. **b)** qRT-PCR expression analysis or Rnf43 and Znrf3 in total liver, hepatocytes, ductal cells (EpCAM+), endothelial cells (CD31+) and macrophages (CD11b+). n=1 mouse. **c)** Schematic representation of Rnf43 and Znrf3 floxed alleles. Grey boxed, exons; red boxes, exons deleted after recombination; yellow triangles, loxP sites. **d)** qRT-PCR liver expression analysis of Rnf43 and Znrf3 in R&Z^flox^, tamoxifen-injected AlbCreERT2-R&Z^del^ and AAV8-Cre-injected R&Z^del^ mice at all time points analysed. Data represent mean ± SD. n=3-5 mice per group. 1W, 1 week; 3M, 3 months; 7M, 7 months; ND, not detected. **e)** qRT-PCR liver expression analysis of Rnf43, Znrf3 and Axin2 in AlbCreERT2-R&Z^flox^ mice with and without tamoxifen. n=1 mouse per condition.

**Supplementary figure 2.**
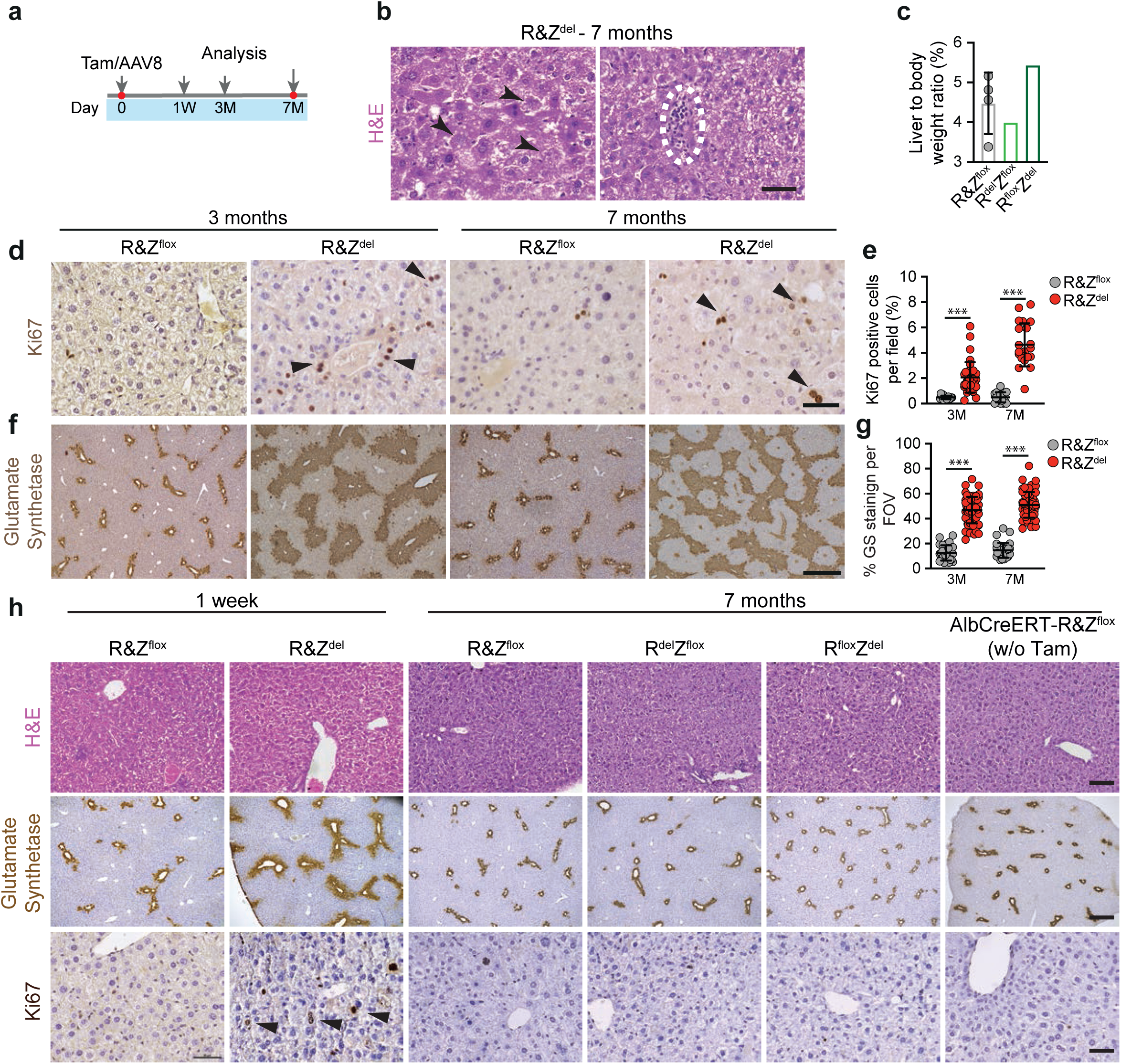
Single deletion of *Rnf43* or *Znrf3* (R&Z) and *AlbCreERT-R&Z*^*flox*^ show normal histology and GS and Ki67 expression and no increase in liver to body weight ratio. **a)** Scheme of experimental design. **b)** Magnification of H&E staining of R&Z^del^ mice after 7 months showing cellular damage (arrowhead) and foci of immunological infiltrate (dotted line). Scale bar is 50μm. **c)** Graph represents the % of liver-to-body weight ratio of mice wild-type (R&Z^flox^) or single deletion (R^del^Z^flox^ and R^flox^Z^del^). Data represent mean +/- SD of n=4 mice for R&Z^flox^ or n=1 for R^del^Z^flox^ and R^flox^Z^del^ livers. **d-g)** *Rnf43* & *Znrf3* liver-specific deletion results in a significant increase in the number of proliferating (Ki67) cells **(d-e)** and the number of glutamine synthetase (GS) positive cells **(f-g)** over time. Representative pictures of immunostainings for Ki67 **(d)** and GS **(f)** on R&Z^flox^ and R&Z^del^ are shown. Scale bar, 500μm (GS) and 50μm (Ki67). Graphs represent the number of Ki67+ hepatocytes (e) and % of GS+ area **(g)** per field-of-view (FOV) are shown. In all cases, data represents mean ± SD from n=2 mice per group. Two-tail t-test was used; ***P<0.001. M, month. **h)** Histopathological analysis of R&Z^flox^ and R&Z^del^ after 1 week deletion, of R^del^Z^flox^ and R^flox^Z^del^ livers and of AlbCreERT-R&Z^flox^ mice w/o tamoxifen. Representative pictures are shown. Scale bar, 100μm (H&E), 500μm (GS) and 50μm (Ki67). Arrowhead, positive Ki67 cells.

**Supplementary figure 3.**
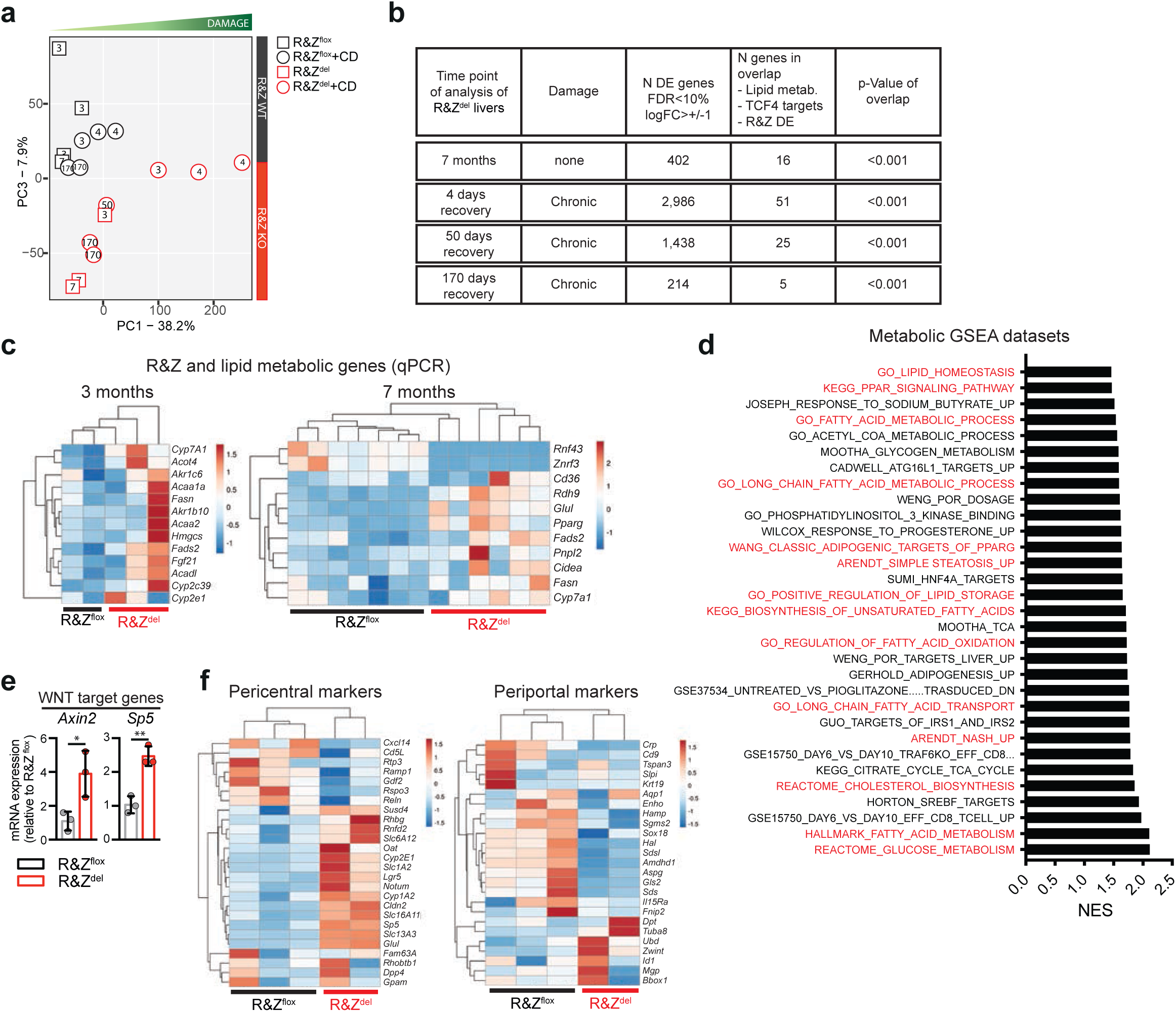
Hyperactivation of the WNT canonical pathway in *Rnf43* & *Znrf3* (R&Z) null hepatocytes results in lipid metabolic changes and a NASH phenotype. **a)** Principal component analysis (PCA) of R&Z^flox^ and R&Z^del^ littermates treated (square) or not (circle) with CCl4 to induce chronic damage (CD) as described in figure 3 and 4. PCA shows RNAseq data (normalized counts) plotted in 2D, using their projections onto two principal components (PC1 and PC3). Each data point represents one sample. Note that PC1 is strongly correlated with the degree of tissue damage, whereas PC3 separates R&Z^flox^ mice from R&Z^del^ mice. **b)** Table showing significance and numbers of overlapped genes between R&Z^del^ DE genes and genes involved in lipid metabolism as well as TCF4 target genes obtained from databases as explained in Fig.2. **c)** Heat maps of qRT-PCR expression analysis (z-scored) of R&Z and lipid metabolic genes at 3 and 7 months after deletion. Experiment was performed twice. **d)** Graphs showing normal enrichment score (NES) for all the significant (FDR<25%; p<0.05) enriched datasets in metabolism and NASH. **e)** Expression analysis (q-RT-PCR) of the indicated Wnt target genes. Graph represents mean±SD of n=3 mice per condition. Circle, individual data point. **f)** Heat-map analysis of the RPKM values (raw z-scored) of liver pericentral and periportal genes indicating that liver metabolic zonation is inverted in Rnf43/Znrf3 mutant livers.

**Supplementary figure 4.**
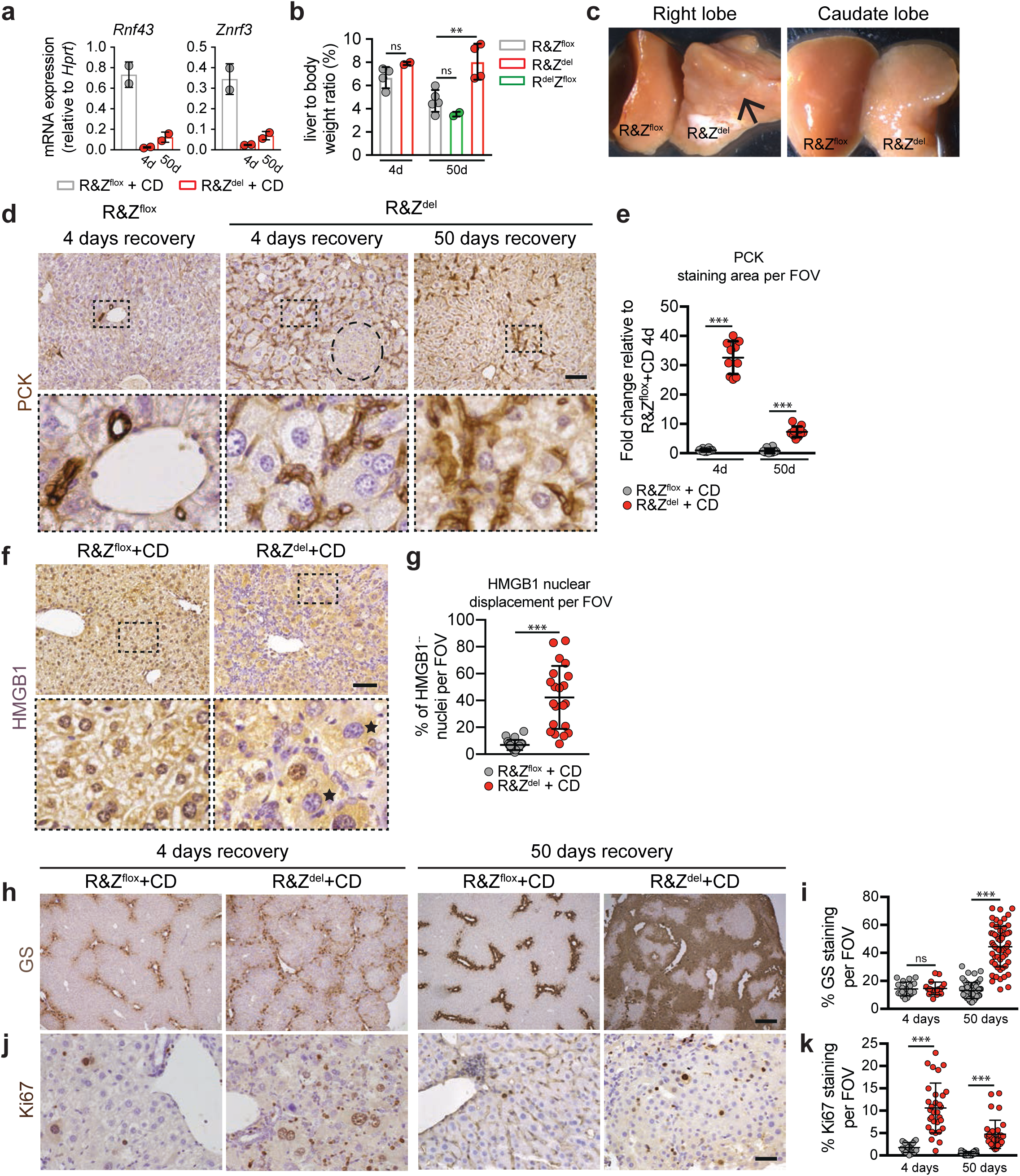
Chronic liver damage after *Rnf43* and *Znrf3* (R&Z) deletion. **a)** qRT-PCR expression analysis or Rnf43 and Znrf3 after chronic damage in R&Z^flox^ and R&Z^del^ mice at 4 and 50 days recovery. Data represent mean +/- SD of n=2 mice per group. **b)** In the settings of chronic liver damage, *Rnf43* & *Znrf3* liver-specific deletion results in hepatomegaly after 50 days recovery. Graph represents the % of liver-to-body weight ratio of mice wild-type (R&Z^flox^) or with double (R&Z^del^) or single (R^del^Z^flox^) deletion of R&Z. Results are presented as mean +/- SD from R&Z^del^-4d and R^del^Z^flox^-50d, n=2; R&Z^flox^-50d, n=5; R&Z^flox^-4d and R&Z^del^-50d, n=4. Two-tail t-test was used; *, p<0.01. **c)** Macroscopic pictures of the right and caudate lobes of R&Z^flox^ and R&Z^del^ mice 50 days in recovery. Arrow points at the irregular/nodular surface of R&Z^del^ livers. **d)** Livers mutated in R&Z show improper tissue regeneration after chronic damage and present ductular reaction as shown by increased PCK immunostaining. Representative pictures are shown. Dashed circle, regenerative nodule. Scale bar is 100μm. **e)** Graphs represent fold change increase PCK staining area. n=2 mice per group, 5 fields per mouse. Data represent mean ± SD; ***P<0.001; two-tail t-test was used. **f)** Representative pictures of the cell damage marker HMGB1 immunostaining. Stars, nuclei negative for HMGB1. Scale bar, 100μm. **g)** Graph shows quantification of HMGB1 nuclear displacement. Data represent mean ± SD of n=2 mice per group, 10 fields of view (FOV) per mouse. **h-k)** Representative picture of immunostaining for GS **(h)** and Ki67 μ of R&Z^flox^ and R&Z^del^ liver sections after chronic damage with respective quantification **(i, k)**. Scale bar, 500μm (GS) and 50μm (Ki67). Data represent mean ± SD of n=2 mice per group, minimum 10 fields per mouse. Two-tail t-test was used; ***P<0.001; ns, not significant.

**Supplementary figure 5.**
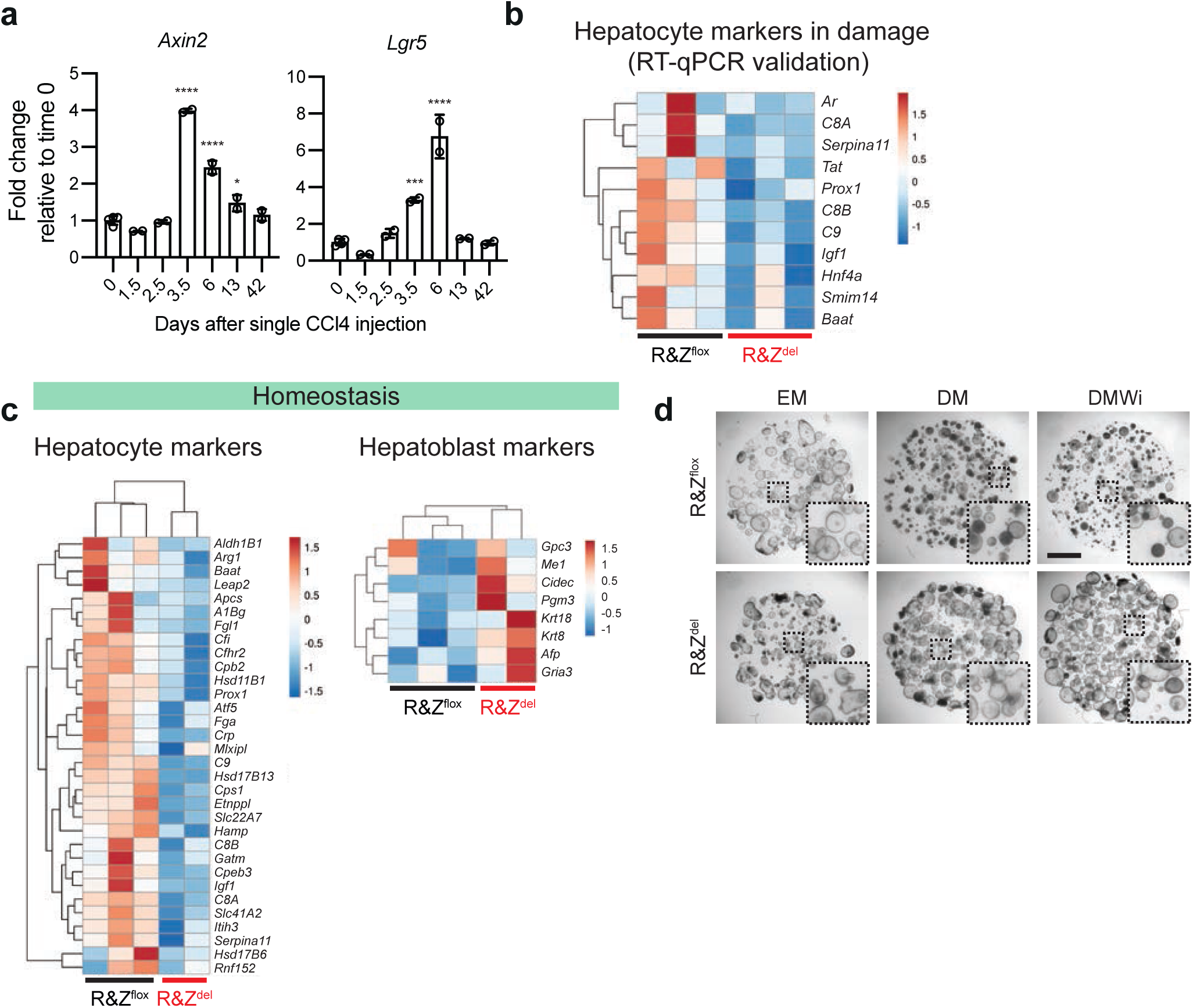
*Rnf43* and *Znrf3* (R&Z) deletion results in impaired maturation of hepatocytes. **a)** qRT-PCR expression analysis of the WNT target genes Lgr5 and Axin2 at several days after a single dose of CCl4. Data represent mean ± SD of 2 mice per condition. **b)** Heat-map analysis of the fold change relative to *Hprt* (raw z-scored) of selected hepatocyte markers. **c)** Heat-map analysis of the RPKM values (raw z-scored) of the hepatocyte and hepatoblasts genes from the GSEA. **d)** Brightfield pictures of organoids from R&Z^flox^ and R&Z^del^ livers in expansion medium (EM), differentiation medium (DM) and differentiation medium with inhibition of the WNT pathway (DMWi). Scale bar is 2mm.

**Supplementary figure 6.**
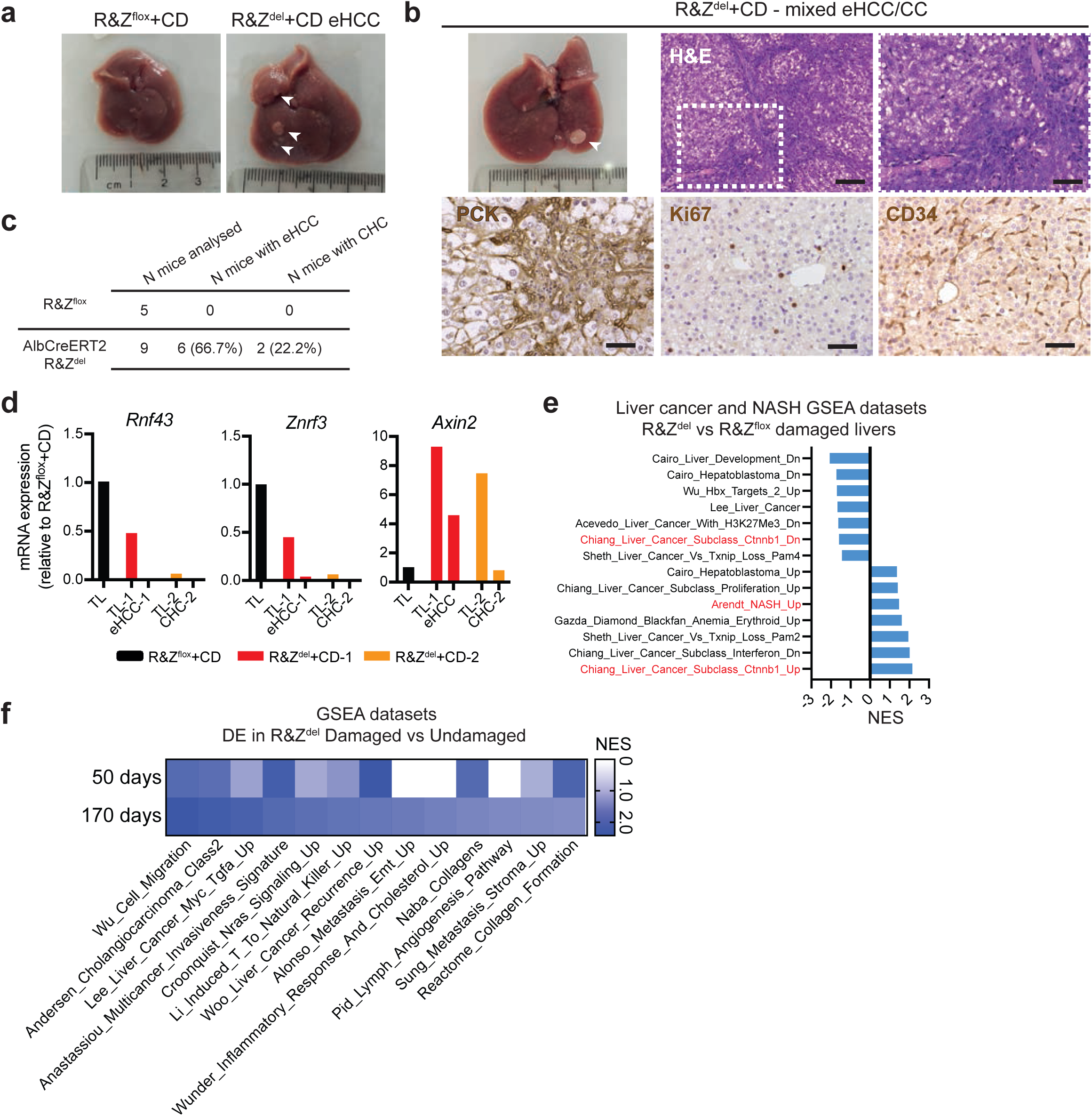
Liver specific *Rnf43* and *Znrf3* (R&Z) deletion leads to hepatocellular carcinoma and tumours with mixed subtype after chronic CCl4 treatment. **a)** Macroscopic pictures of healthy R&Z^flox^+CD liver and eHCC lesion of R&Z^del^+CD liver. **b)** Brightfield picture and representative pictures of H&E staining and PCK, Ki67 and CD34 immunostaining of a mixed subtype neoplastic lesion (CHC). Scale bar, 100μm (H&E left panel); 50μm (PCK, Ki67, CD34 and H&E right panel). **c)** Table showing number of animals involved in the study and penetrance of eHCC and CHC. **d)** Expression pattern (qRT-PCR) of *Rnf43, Znrf3 and Axin2* in one eHCC and one CHC lesion. TL, total liver. eHCC or CHC, microdissected lesion **e)** Graph showing normal enrichment score (NES) for all liver cancer GSEA datasets significantly enriched (FDR<25%, pV<0.05). **f)** Heatmap showing normal enrichment score (NES) for selected GSEA datasets R&Z^del^ damaged livers against R&Z^del^ undamaged livers at 50 and 170 days of recovery. Blue, pV<0.05. White, pV>0.05.

**Supplementary figure 7.**
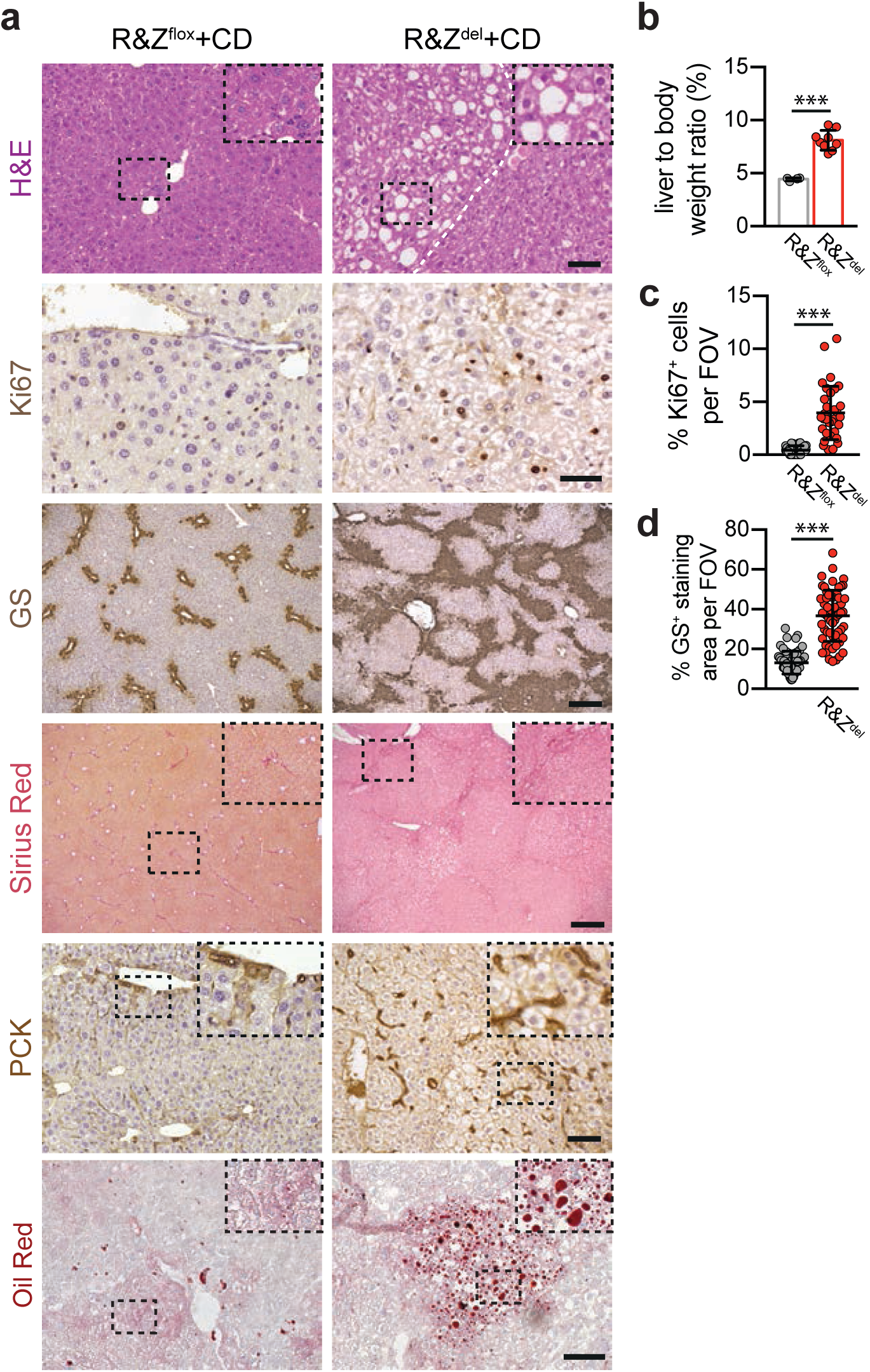
Chronic liver damage after long-term R&Z deletion. **a)** Representative pictures of H&E, Sirius red and Oil Red O staining and immunostaining for Ki67, GS and PCK of chronically damaged R&Z^flox^ and R&Z^del^ liver sections after 170 days of recovery. Dotted white lines mark nodule borders. Scale bar, 200μm (H&E, GS, Sirius Red); 100μm (Oil red, PCK), 50μm (Ki67). **b)** Graph showing differences in the percentage of liver to body weight ratio. n=3 mice (R&Z^flox^); 9 mice (R&Z^del^). Data represent mean ± SD of at least 5 mice per group. Two-tail t-test was used; ***P<0.001. **c-d)** Quantification of Ki67 positive hepatocytes per field of view **(c)** and of GS positive staining localized around central veins **(d)**. Data represent mean ± SD of n=3 mice per group. Two-tail t-test was used; ***P<0.001.

